# Characterizing Tyrosine Ring Flips in Proteins by ^19^F NMR

**DOI:** 10.1101/2024.12.29.630691

**Authors:** Meng Lu, Wenkai Zhu, Guohua Xu, Jiawen Chen, Qiong Wu, Xiaoli Liu, Mingjuan Yang, Ling Jiang, Minghui Yang, Maili Liu, Conggang Li

**Author notes:** To whom correspondence should be addressed. (Conggang Li) (Guohua Xu). These authors contributed equally to this work (Meng Lu, Wenkai Zhu).

## Abstract

Aromatic ring-flip dynamics are hallmarks of concerted protein “breathing” motions that are essential for biological function. Ring flips occur on a broad range of timescales (ns–s) and have been primarily studied by NMR spectroscopy, typically requiring expensive isotope labeling of aromatic side chains, thereby limiting such studies to a few proteins. Here, we report two novel di-fluorotyrosine probes, 3,5-F_2_Y and 2,6-F_2_Y, that can be incorporated into proteins cost-effectively for characterizing and modulating tyrosine ring-flip dynamics. We show that ^19^F rotating-frame (*R*_1*ρ*_) relaxation dispersion is powerful for quantifying ring flips on the µs–ms timescale. Importantly, the tyrosine ring-flip rates (*k*_flip_) for 3,5-F_2_Y in GB1 and HPr are comparable to those measured by ^1^H and ^13^C NMR studies, validating 3,5-F_2_Y as a largely non-perturbing, native-like ring-flip probe. In contrast, 2,6-F_2_Y acts as an effective “brake” on ring flips, enabling direct visualization and quantitative characterization of previously undetected tyrosine ring flips in ubiquitin via ^19^F lineshape analysis. Furthermore, we use 2,6-F_2_Y to assess the environmental effects on ring-flip dynamics in crowding reagents and in living *X. laevis* oocytes. Collectively, our study opens a new avenue for measuring and modulating ring-flip dynamics *in vitro* and in living cells.

**TOC:** 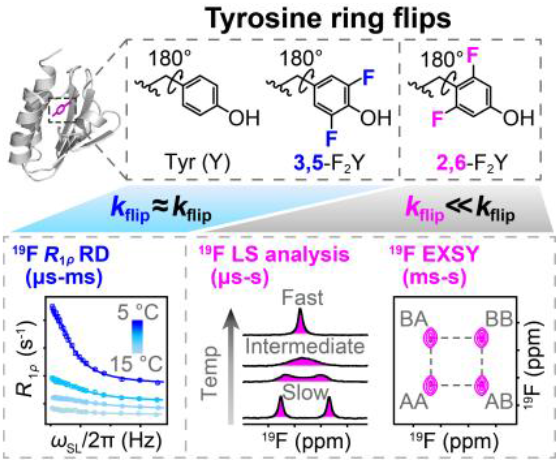

## Introduction

Aromatic residues are frequently found in the hydrophobic core of folded proteins and involved in π-π, CH-π, and cation-π interactions.^[1-5]^ The aromatic rings of phenylalanine (Phe) and tyrosine (Tyr) undergo ring flips, *i*.*e*., 180° rotations of the *χ*_2_ dihedral angle around the C_β_-C_γ_-C_ζ_ axis (Figure 1a). The flip motions of buried aromatic rings are enabled by transient cavities formed by surrounding residues, thus serving as sensitive reporters of concerted protein “breathing” motions that are essential for biological functions.^[6-7]^ Recent studies have further highlighted the importance of ring flips in mediating protein structural transitions between ground and high-energy states and essential conformational rearrangements during receptor activation.^[8-9]^ Capturing ring-flip motions therefore reveals functionally relevant protein dynamics and uncovers detailed molecular mechanisms underlying protein breathing, which can be further elucidated from the structural and energetic properties of the ring flipping transition state, *i*.*e*., the activation enthalpy (*ΔH*^‡^), activation entropy (*ΔS*^‡^), activation volume (*ΔV*^‡^), and isothermal volume compressibility (*Δκ*^‡^).^[10-20]^ Since the pioneering NMR observations of ring flips in the 1970s,^[21-25]^ various solution NMR techniques have been employed for characterizing ring-flip dynamics across µs–s timescales, including ^1^H-^1^H exchange spectroscopy (EXSY),^[14, 17, 26]^ longitudinal exchange spectroscopy,^[27]^ cross-saturation,^[23] 1^H lineshape (LS) analysis,^[16, 18-20, 22-23] 1^H/^13^C CPMG and on/off-resonance rotating-frame (*R*_1*ρ*_) relaxation dispersion (RD),^[11-13, 28-31]^ and ^13^C/^19^F spin relaxation (*R*_1_/*R*_2_/NOE) ^[32-33]^. In addition, computational studies have provided atomistic insights into the ring-flip mechanisms and sequential events accompanied by ring flips.^[8, 34-37]^ Nevertheless, ring-flip studies have so far been limited to a few proteins, including BPTI,^[15, 17-18, 20, 22, 29-30, 38-39]^ Cyt *c*,^[19, 23, 26]^ GB1,^[11-13, 28]^ BcX,^[27]^ HPr,^[16]^ FKBP12,^[14]^ alkaline phosphatase^[32]^, ubiquitin,^[33, 35]^ TET2,^[40]^ amyloid fibrils,^[41]^ and the fd bacteriophage coat protein,^[42]^ presumably because of the expensive specific isotope labeling, complicated spectral assignments, and, more importantly, the lack of sensitive and universal ring-flipping probes which can access a wide range of timescales (ns–s) on which ring flips occur.

**Figure 1.**
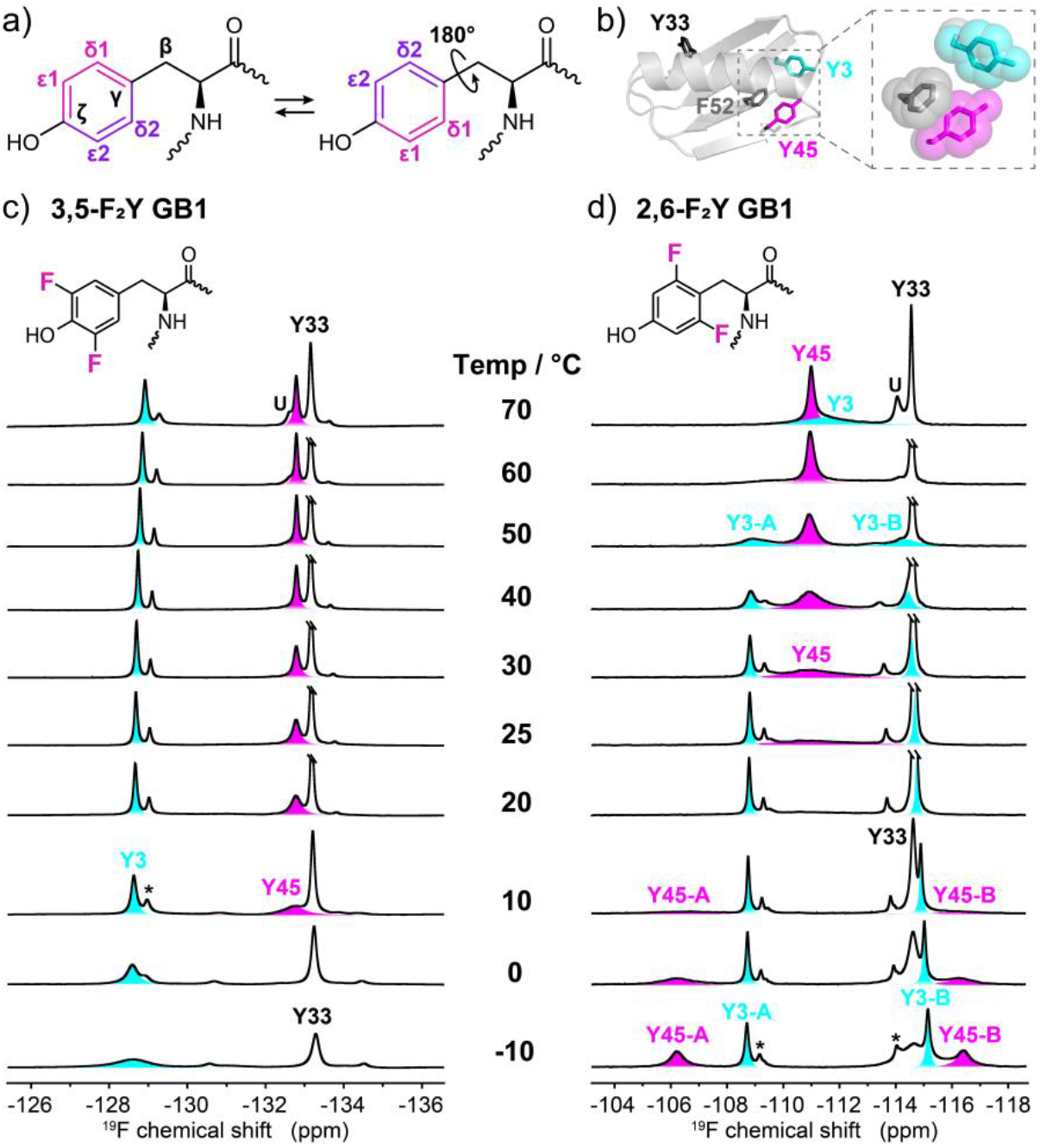
Temperature-dependent ^19^F spectra of fluorinated GB1. a) Schematic illustration of a tyrosine residue undergoing 180° ring flips around the C_β_-C_γ_-C_ζ_ axis. b) Cartoon representation of GB1 (PDB id: 2QMT), with side chains of Y3 (cyan), Y45 (magenta), Y33 (black), and F52 (gray) shown as sticks. The aromatic cluster formed by Y3, F52, and Y45 is shown enlarged in the dashed rectangle. ^19^F spectra of c) 3,5-F_2_Y and d) 2,6-F_2_Y GB1 recorded from -10 °C to 70 °C. Peaks for Y3 and Y45 are shaded in cyan and magenta, respectively, to highlight temperature-dependent ^19^F lineshape changes because of the ring-flip motion. Chemical structures of 3,5-F_2_Y and 2,6-F_2_Y are shown above each panel. The signals arising from incomplete fluorine incorporation are labeled by asterisks.

Fluorine-19 (^19^F) is an attractive probe due to its 100% natural abundance, high sensitivity (83% relative to ^1^H), broad chemical shift dispersion and acute responsiveness to local environments.^[43-45]^ Its absence in biological systems further establishes ^19^F as a unique reporter for in-cell studies.^[46-47]^ Among ^19^F-labeled biomolecular probes, fluorinated aromatic amino acids have emerged as particularly powerful tools for interrogating protein structure, dynamics, and interactions.^[48-54]^ Hull and Sykes pioneered the qualitative assessment of tyrosine ring-flip timescales using 3-fluorotyrosine in alkaline phosphatase in 1975,^[32]^ yet subsequent applications of ^19^F NMR to aromatic ring-flip dynamics have remained scarce. Fluorinated tyrosine analogs offer several advantages for characterizing ring flips. First, they can be efficiently incorporated into proteins cost-effectively via *E. coli* expression.^[55-56]^ Second, the ^19^F chemical shift difference between two exchanging states (*Δω*_AB_) is generally larger than that of ^1^H, ^13^C and ^15^N, frequently placing ring-flip motions in a slower exchange regime which is ideal for NMR relaxation dispersion studies.^[57-58]^ Third, one-dimensional ^19^F detection enables direct observation of two equally populated states (ring-flipping states) in the intermediate-to-fast exchange regime where the exchange contribution *R*_ex_ (≈ *p*_A_>*p*_B_>Δω^2^ ∕ *k*_ex_) reaches the maximum, nearly impossible by ^1^H and ^13^C relaxation studies due to the excessive line-broadening. Finally, ^19^F aromatic amino acids enable characterization of ring-flip dynamics in living cells, where ^1^H and ^13^C based NMR studies are infeasible due to severe cellular background.

Here we present a pair of symmetric ring-flip ^19^F probes, 3,5-F_2_Y and 2,6-F_2_Y, in which the H^*ε*1, *ε*2^ and H^*δ*1, *δ*2^ of the tyrosine aromatic ring are substituted with ^19^F atoms, respectively. Both 3,5-F_2_Y and 2,6-F_2_Y can be readily incorporated into proteins via natural biosynthesis in *E. coli*. We monitored ring-flip dynamics by collecting the ^19^F spectra at varying temperatures. Accurate ring-flip rates (*k*_flip_) and activation parameters (*ΔH*^‡^ and *ΔS*^‡^) were obtained by quantitative ^19^F lineshape (LS) analysis over a broad temperature range, in which ring flips traverse from slow- to fast-exchange regimes. We show that ^19^F rotating-frame (*R*_1*ρ*_) relaxation dispersion (RD) experiments are powerful for studying ring-flip dynamics on the µs–ms timescale when a single averaged peak is observed, whereas ^19^F-^19^F exchange spectroscopy is essential for determining *k*_flip_ rates in the slow exchange (ms–s) timescale. The joint use of LS analysis, EXSY, and *R*_1*ρ*_ RD thus provides a robust and reliable approach for quantifying ring-flip dynamics on the µs–s timescale. The *k*_flip_ values for 3,5-F_2_Y are similar to those of the native tyrosines, whereas 2,6-F_2_Y significantly slows ring flips in GB1, HPr, and Ub. The hampered ring flips of 2,6-F_2_Y in different proteins suggest that it generally slows ring flips by raising the activation barrier, through ground-state stabilization and/or transition-state destabilization. Notably, using 2,6-F_2_Y, we directly observed the Y59 ring flip in Ub, which previous studies indicated does not occur below the transition temperature.^[33]^ Given that ring flips are widely recognized as a hallmark of protein “breathing” motions, 2,6-F_2_Y now opens a new avenue for unveiling their roles in protein “breathing” motions and biological functions. Moreover, we show that ring-flip rates can be accurately quantified in *X. laevis* oocytes, enabling the evaluation of cellular environmental effects on protein ring flips. Taken together, our work establishes an efficient and cost-effective strategy that employs two sensitive ^19^F probes, 3,5-F_2_Y and 2,6-F_2_Y, to visualize and modulate tyrosine ring-flip dynamics both *in vitro* and in cells.

## Results and Discussion

### Incorporation of 3,5-F_2_Y and 2,6-F_2_Y into GB1

Aromatic ring-flip motion is characterized by the 180° rotation of the *χ*_2_ dihedral angle (around the C_*β*_-C_*γ*_ axis), producing two symmetry-equivalent states (Figure 1a). To probe ring-flip motions, we enzymatically synthesized two symmetrical di-fluorinated tyrosines, 3,5-F_2_Y and 2,6-F_2_Y (Figure S1a).^[59]^ In both cases, the two ring-flipping states will be equally populated, which facilitates quantitative analysis and distinction of ring flips from other conformational exchange processes.^[28]^ The identification and purity of fluorinated tyrosines were confirmed by ESI-MS and ^19^F NMR (Figure S1b–d). Both 3,5-F_2_Y and 2,6-F_2_Y were successfully incorporated into GB1 via *E. coli* expression and incorporation efficiencies were determined by ESI-MS (Figure S1e–f), and the ^19^F peaks were assigned by site-directed mutagenesis (Figure S2). Fluorination minimally perturbed the overall structure of GB1, as evidenced by ^1^H-^15^N HSQC spectra (Figure S3a). Importantly, circular dichroism (CD) spectra and ^15^N *R*_1_/*R*_2_ relaxation data suggest negligible fluorination effects on protein thermal stability and backbone dynamics, respectively (Figure S3b–c).

We acquired 1D ^19^F spectra of 3,5-F_2_Y and 2,6-F_2_Y GB1 from -10 °C to 70 °C, using supercooled water to achieve subzero temperatures (Figure 1c–d).^[17]^ For 2,6-F_2_Y labeled Y3 and Y45, two distinct ^19^F resonances with a 1:1 population ratio are observed between -10 °C and 10 °C, which coalesce into a broad peak and then sharpen at higher temperatures, directly reflecting the ring-flip motions of Y3 and Y45 in GB1. By contrast, a single averaged peak is observed for the Y3 and Y45 in 3,5-F_2_Y GB1. This difference indicates that ring flips of the same tyrosine, using 2,6-F_2_Y versus 3,5-F_2_Y probe, occur on different NMR timescales, which relies on the relative difference of ring-flip rates (*k*_flip_) and ^19^F chemical shift differences (*Δω*_AB_) at the δ (2,6) and ε (3,5) positions. For Y33, however, the ^19^F peaks in both 3,5-/2,6-F_2_Y GB1 show minimal temperature dependence, indicating a much faster *k*_flip_ and/or a small *Δω*_AB_, consistent with the previous reports.^[11, 13]^ Collectively, those observations demonstrate that both 3,5- and 2,6-F_2_Y are sensitive probes for detecting ring flips in GB1, enabling quantitative determination of ring-flip rates by ^19^F lineshape analysis, ^19^F-^19^F exchange spectroscopy (EXSY), and *R*_1*ρ*_ relaxation dispersion (RD) experiments (see below).

### Quantifying tyrosine ring-flip rates in 3,5-F_2_Y and 2,6-F_2_Y GB1

In 3,5-F_2_Y GB1, significant ^19^F line broadening was observed for Y3 and Y45 over the studied temperature range (Figure 1c), indicating µs–ms ring-flip motions, as reported previously.^[11, 13]^ NMR relaxation dispersion (RD) experiments are ideal for probing protein dynamics on the µs–ms timescale. Given the large ^19^F line widths, we performed on-resonance ^19^F rotating-frame (*R*_1*ρ*_) RD experiments, which can sample arbitrary spin-lock frequencies regardless of short ^19^F T_2_ time. Notably, ^19^F *R*_1*ρ*_ RD is little affected by ^3^J_HF_ couplings (∼ 10 Hz), while ^3^J_HH_ couplings are detrimental for ^1^H (H^*ε*1, *ε*2^) *R*_1*ρ*_ RD studies and deuteration of vicinal protons is essential to record artifact-free RD profiles by using ^2^H/^13^C ɑ-ketoacid precursors.^[11]^ High-quality ^19^F *R*_1*ρ*_ RD profiles for 3,5-F_2_Y3 and 3,5-F_2_Y45 were recorded over their respective optimal temperature ranges, despite substantial peak broadening (*R*_2_ > 150 s^-1^) (Figure 2a–b, Table 1). Gratifyingly, large relaxation dispersions (*R*_ex_ ∼ 60–130 s^-1^) for both 3,5-F_2_Y3 and 3,5-F_2_Y45 permit accurate determination of *k*_flip_ (Figure 2a–b). The ^19^F *R*_1*ρ*_ RD profiles for solvent-exposed Y33 is essentially flat, indicating its ring flip occurs on a faster NMR timescale (Figure S4). The *R*_1*ρ*_ RD-derived ^19^F chemical shift differences *Δω*_AB_ for Y3 is 1.1 ± 0.1 ppm, roughly twice the value reported for ^1^H (0.5 ppm),^[11]^ which results in a 4-fold larger dispersion for same *k*_flip_ rates. The activation enthalpies and entropies for 3,5-F_2_Y3/Y45 ring flips were derived from *k*_flip_ using Eyring equation (Figure 2c, Table 3).^[60]^

**Table 1.** *k*_flip_ (10^3^ s^-1^) determined by ^19^F *R*_1*ρ*_ RD for 3,5-F_2_Y labeled proteins.

| T (°C) | GB1, Y3 | GB1, Y45 | HPr, Y6 | Ub, Y59 |
| --- | --- | --- | --- | --- |
| 30.0 | | $77 \pm 7$ | | |
| 25.0 | | $51 \pm 5$ | | |
| 20.0 | | $34 \pm 3$ | | |
| 15.0 | | | $10.6 \pm 0.8$ | $123 \pm 27$ |
| 12.5 | $43 \pm 2$ | | $9.2 \pm 0.3$ | |
| 10.0 | $35 \pm 2$ | | $7.4 \pm 0.3$ | $90 \pm 20$ |
| 7.5 | $28 \pm 2$ | | $5.5 \pm 0.5$ | |
| 5.0 | $22 \pm 1$ | | $4.1 \pm 1.0$ | $60 \pm 14$ |

**Figure 2.**
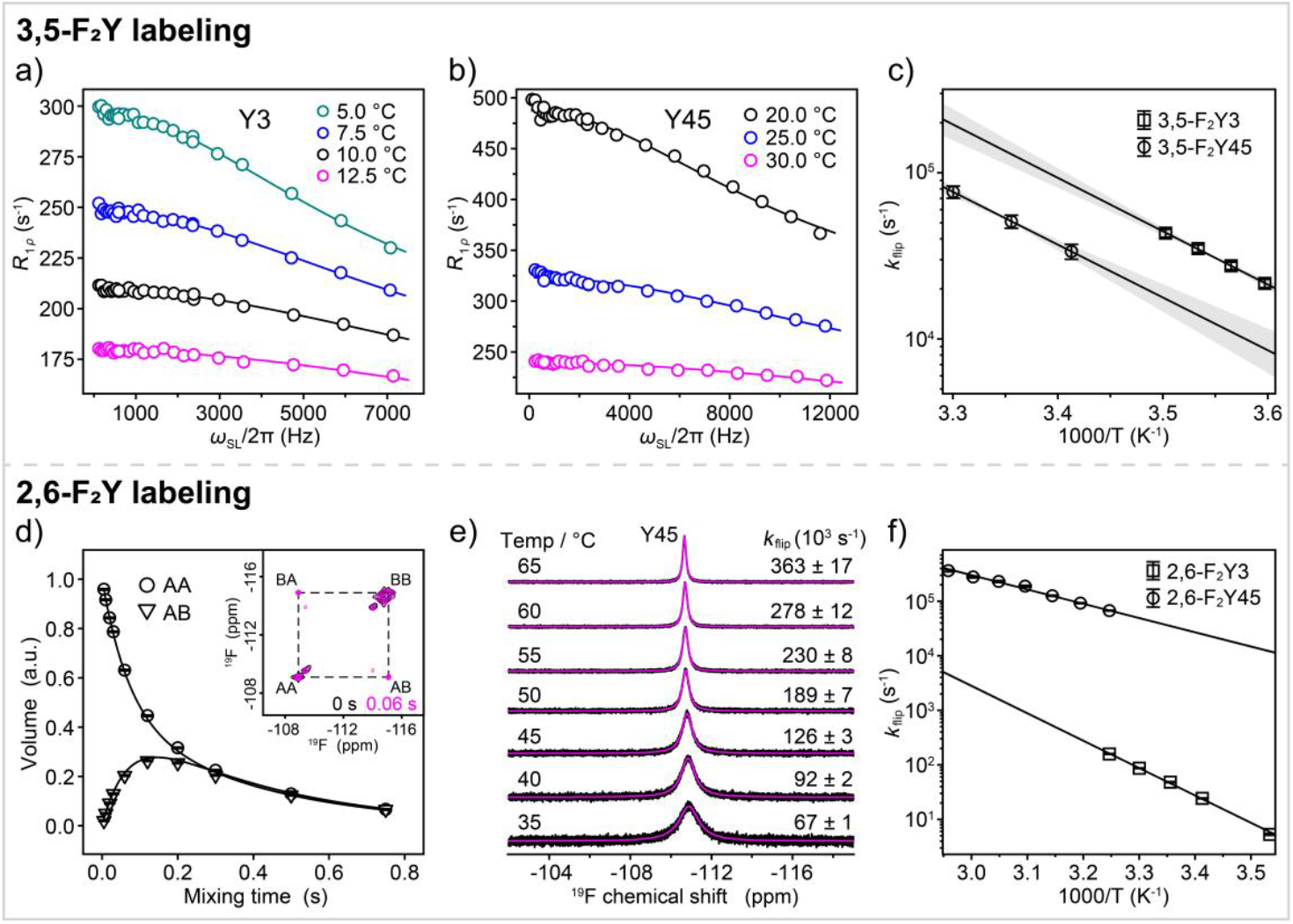
Ring-flip dynamics of 3,5-/2,6-F_2_Y GB1. On-resonance ^19^F *R*_1*ρ*_ RD profiles recorded at 16.4 T for 3,5-F_2_Y GB1 are shown for a) 3,5-F_2_Y3 at 5 °C (cyan), 7.5 °C (blue), 10 °C (black), and 12.5 °C (magenta) and b) 3,5-F_2_Y45 at 20 °C (black), 25 °C (blue), and 30 °C (magenta). The ^19^F *R*_1*ρ*_ RD data were fitted by numerical propagation of the Bloch-McConnell equations with populations *p*_A_ = *p*_B_ = 0.5 and Δ*ω*_AB_ as a free parameter. The derived *k*_flip_ values are given in Table 1. The *R*_2_ errors were estimated from replicate experiments at two spin-lock fields (*ω*_SL_/2π = 500 and 2000 Hz). c) Temperature dependence of the ring-flip rates (*k*_flip_) derived from *R*_1*ρ*_ RD in a) for Y3 and in b) for Y45, fitted to the Eyring equation to yield activation enthalpy *ΔH*^‡^ and entropy *ΔS*^‡^, listed in Table 3. d) ^19^F-^19^F EXSY experiment for measuring the ring-flipping rate of Y3 in 2,6-F_2_Y GB1. Auto (AA) and exchange (AB) peak volumes were fitted as a function of mixing times at 10 °C. The errors in peak volumes were estimated from the spectrum noise. Inset shows EXSY spectra with a mixing time of 0 s (black) and 0.06 s (magenta) with auto (AA and BB) and exchange (AB and BA) peaks originating in the A and B state labeled by the dashed lines. e) ^19^F lineshape analysis of Y45 in 2,6-F_2_Y GB1 from 35 °C to 65 °C. Experimental ^19^F NMR spectra for 2,6-F_2_Y GB1 (Figure 1d) were deconvoluted (black) and subsequently fitted (magenta) as described in “Materials and Methods”, with *k*_flip_ values indicated for each temperature. f) Temperature dependence of the ring-flip rates (*k*_flip_) derived from EXSY in d) for Y3 and from ^19^F LS analysis in e) for Y45, fitted to the Eyring equation to derive activation parameters (*ΔH*^‡^ and *ΔS*^‡^), listed in Table 3.

We further investigated Y3 and Y45 ring-flip dynamics in 2,6-F_2_Y GB1, showing two ring-flipping states in slow exchange on NMR timescales (Figure 1d). The *k*_flip_ for 2,6-F_2_Y3 was determined by EXSY from 10 to 35°C (Figure 2d, Table 2). Remarkably, 2,6-F_2_Y incorporation slows down Y3 ring flip by three magnitude orders, from 35.0 × 10^3^ s^-1^ for 3,5-F_2_Y to 5.8 s^-1^ for 2,6-F_2_Y at 10 °C (Table 1–2). Both the activation enthalpy and entropy for Y3 are significantly larger in 2,6-F_2_Y GB1 (*ΔH*^‡^ = 94 ± 1 kJ·mol^-1^, *ΔS*^‡^ = 101 ± 3 J·mol^-1^ K^-1^) than in 3,5-F_2_Y GB1 (*ΔH*^‡^ = 59 ± 7 kJ·mol^-1^, *ΔS*^‡^ = 51 ± 24 J·mol^-1^ K^-1^) (Figure 2f, Table 3), suggesting that 2,6-F_2_Y incorporation stabilizes the ground state and/or destabilizes the transition state of Y3. Considering that fluorine atoms at 2,6-positions are closer to the backbone atoms than those at 3,5-positions (Figure 1c–d), we hypothesize that the hampering effects partially arises from the intra-residue repulsion. In addition, we performed molecular simulations of ring flips for 3,5-/2,6-F_2_Y and non-fluorinated AYA tripeptide, where interactions with surrounding residues are absent. We found the activation barrier for the ring-flip motion is 15 kJ/mol higher for 2,6-F_2_Y than for 3,5-F_2_Y and natural tyrosine, which are essentially the same (Figure S5). Although intra-residue repulsive interactions may hamper ring flips in 2,6-F_2_Ys, inter-residue interactions also play important roles in modulating ring flips, and their overall effect determines the *k*_flip_ of 2,6-F_2_Ys. To corroborate this, we found that *k*_flip_ for 2,6-F_2_Y45 determined from ^19^F lineshape analysis is only slightly lower than that of 3,5-F_2_Y45 at 30 °C (Figure 2e, Table 2), as 2,6-F_2_Y45 can potentially alter aromatic π-π stacking interactions with F52 (Figure 1b), thereby mitigating the hampering effects. Importantly, when the ring flip of 2,6-F_2_Y3 is almost abolished at 10 °C (*k*_flip_ = 5.8 s^-1^), 2,6-F_2_Y45 still flips frequently (*k*_flip_ = 14 × 10^3^ s^-1^), supporting independent ring flips of Y3 and Y45 despite their tight packing within the aromatic cluster.

**Table 2.** *k*_flip_ (s^-1^) determined by ^19^F lineshape analysis for 2,6-F_2_Y labeled proteins.

| T ($^{\circ}\text{C}$) | GB1, Y3 | GB1, Y45 | Y64F HPr, Y6 | Ub, Y59 |
| --- | --- | --- | --- | --- |
| 65 | | $(363 \pm 17) \times 10^3$ | | |
| 60 | | $(278 \pm 12) \times 10^3$ | | |
| 55 | | $(230 \pm 8) \times 10^3$ | $13246 \pm 821$ | |
| 50 | | $(189 \pm 7) \times 10^3$ | $6763 \pm 158$ | |
| 45 | | $(126 \pm 3) \times 10^3$ | $3256 \pm 35$ | |
| 40 | | $(92 \pm 2) \times 10^3$ | $1112 \pm 2$ | $(31 \pm 5) \times 10^3$ |
| 35 | $158.2 \pm 2.5^{[a]}$ | $(67 \pm 1) \times 10^3$ | $500 \pm 12$ | |
| 30 | $86.6 \pm 1.1^{[a]}$ | $49 \times 10^3^{[b]}$ | | $(18 \pm 2) \times 10^3$ |
| | | | | $(25 \pm 1) \times 10^3^{[c]}$ |
| 25 | $48.2 \pm 1.1^{[a]}$ | | $61.2 \pm 1.7^{[a]}$ | $(14 \pm 1) \times 10^3$ |
| 20 | $24.3 \pm 0.4^{[a]}$ | | | $(11 \pm 1) \times 10^3$ |
| 10 | $5.8 \pm 0.2^{[a]}$ | $14 \times 10^3^{[b]}$ | $3.4 \pm 0.1^{[a]}$ | $5336 \pm 219$ |
| 0 | | | | $2368 \pm 49$ |
| -5 | | | | $1336 \pm 46$ |
| -8 | | | | $1017 \pm 64$ |

[a] The $k_{\text{flip}}$ for 2,6- $\text{F}_2\text{Y}$ 3 in GB1 was determined by $^{19}\text{F}$ - $^{19}\text{F}$ EXSY. [b] The $k_{\text{flip}}$ for 2,6- $\text{F}_2\text{Y}$ 45 in GB1 at 10 $^{\circ}\text{C}$ and 30 $^{\circ}\text{C}$ was extrapolated using the Eyring equations. [c] The $k_{\text{flip}}$ for 2,6- $\text{F}_2\text{Y}$ 59 in Ub was determined by $^{19}\text{F}$ $R_{1\rho}$ RD.

**Table 3.** Activation parameters of tyrosine ring flips fitted by the Eyring equation.

| Protein | Residue | $\Delta H^\ddagger$<br>(kJ mol<sup>-1</sup>) | $\Delta S^\ddagger$<br>(J mol<sup>-1</sup> K<sup>-1</sup>) |
| --- | --- | --- | --- |
| GB1 | 3,5-F<sub>2</sub>Y3 | 59 ± 7 | 51 ± 24 |
|  | 2,6-F<sub>2</sub>Y3 | 94 ± 1 | 101 ± 3 |
|  | 3,5-F<sub>2</sub>Y45 | 58 ± 10 | 42 ± 33 |
|  | 2,6-F<sub>2</sub>Y45 | 48 ± 1 | 2 ± 3 |
| HPr | 3,5-F<sub>2</sub>Y6 | 58 ± 8 | 35 ± 29 |
|  | 2,6-F<sub>2</sub>Y6<sup>[a]</sup> | 145 ± 1 | 277 ± 2 |
| Ub | 2,6-F<sub>2</sub>Y59 | 49 ± 1 | -1 ± 4 |

[a] Y64F HPr was used for <sup>19</sup>F lineshape analysis.

We next compared *k*_flip_ values obtained from ^19^F *R*_1*ρ*_ RD for 3,5-F_2_Y with those reported for native tyrosines in GB1.^[11, 13]^ MD simulations have shown that the activation free energies for ring flips of 3,5-F_2_Y and native tyrosines in the tripeptide AYA are similar. Nevertheless, we found that the *k*_flip_ for 3,5-F_2_Y3 and 3,5-F_2_Y45 are approximately 8- and 5-fold larger, respectively, than those previously reported from ^1^H/^13^C *R*_1*ρ*_ RD studies (Table S2). These discrepancies may arise from differences in experimental conditions, the quality of RD profiles, and fluorination effects of 3,5-F_2_Y incorporation, which can slightly perturb inter-residue interactions in the aromatic cluster, as discussed above. Both *ΔH*^‡^ and *ΔS*^‡^ for 3,5-F_2_Y are lower than those previously reported for native tyrosine, whereas the activation *ΔG*^‡^ values are in reasonable agreement (Table 3). This likely reflects the strong correlation between *ΔH*^‡^ and *ΔS*^‡^, which depends on the accuracy of fitted *k*_flip_. Notably, large discrepancies in *ΔH*^‡^ and *ΔS*^‡^ for F52 in GB1 have been reported in different *R*_1*ρ*_ RD studies.^[12-13]^

### Characterizing tyrosine ring-flip dynamics in 3,5-F_2_Y and 2,6-F_2_Y HPr

To further validate 3,5-F_2_Y as a robust, native-like ring-flip probe and to assess whether 2,6-F_2_Y generally hampers the ring-flip motions, we incorporated them into HPr (Figure S6), for which activation parameters have been reported previously.^[16]^ As described above, we confirmed that the fluorination minimally perturbs protein structure, stability and backbone dynamics (Figure S7) and obtained ^19^F peak assignments by site-directed mutagenesis (Figure S8). At 5–20 °C, a single averaged peak was observed for 3,5-F_2_Y HPr (Figure 3a). The dramatic broadening 3,5-F_2_Y6 peak at 5.0 °C (135 Hz) relative to 10.0 °C (82 Hz) clearly indicates the ring-flip motion on the µs–ms timescale, as the rotational correlation time (*τ*_c_) of HPr is increased by only 20% calculated by *HydroPro*.^[61]^ We therefore recorded on-resonance ^19^F *R*_1*ρ*_ RD data for 3,5-F_2_Y HPr at 5.0 °C (black), 7.5 °C (magenta), 10.0 °C (teal), 12.5 °C (blue), and 15.0 °C (orange) (Figure 3b, Table 1). The dispersion amplitude decreases significantly from 5 °C to 15 °C, indicating increasing *k*_flip_ with temperature, similar to the ^19^F *R*_1*ρ*_ RD profiles of 3,5-F_2_Y GB1. We found that ^19^F *R*_1*ρ*_ RD profiles at the five temperatures need to be fitted individually, otherwise the reduced *χ*^2^ becomes significantly higher 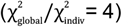 when fitted globally assuming a common *Δω*_AB_, suggesting different ^19^F chemical shift temperature coefficients for the two ring-flipping states. We further confirmed this by recording ^19^F *R*_1*ρ*_ RD data at two magnetic fields (14.1 T and 16.4 T) for more robust fitting of *Δω*_AB_ and *k*_ex_ (Figure S9). The determined *Δω*_AB_ values vary linearly with temperature, and *k*_flip_ exhibits the typical temperature-dependence described by Eyring equation (Figure 3c), in good agreement with reported *k*_flip_ from ^1^H LS analysis (Table S2). However, the resulting activation enthalpy *ΔH*^‡^ (58 ± 8 kJ·mol^-1^) is lower than that determined from ^1^H LS analysis (87 ± 17 kJ·mol^-1^), while the activation entropies (*ΔS*^‡^) are similar (35 ± 29 *v*.*s*. 16 ± 3 J·mol^-1^ K^-1^).^[16]^ Our Monte Carlo simulations show that *k*_flip_ positively correlates with *Δω*_F, AB_ in the fast-exchange regime during lineshape fitting (Figure S10). Assuming a common *Δω*_F, AB_ over the studied temperatures can therefore either overestimate or underestimate the true *k*_flip_. Thus, determining *Δω*_AB_ separately at each temperature is essential for obtaining reliable activation parameters.

**Figure 3.**
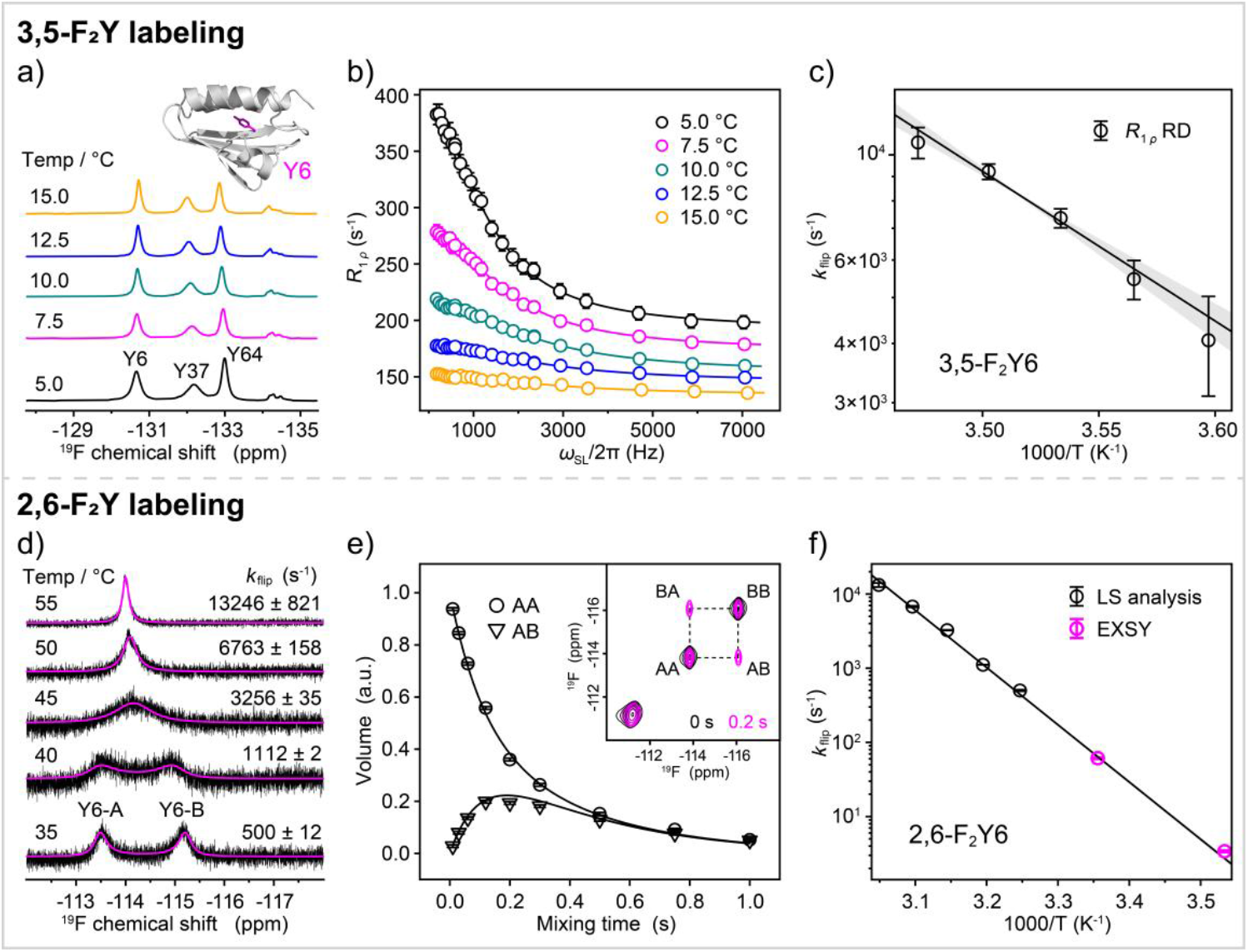
Ring-flip dynamics of Y6 in 3,5-F_2_Y HPr and 2,6-F_2_Y Y64F HPr. a) Experimental ^19^F spectra recorded at 5 °C (black), 7.5 °C (magenta), 10.0 °C (teal), 12.5 °C (blue), and 15 °C (orange) for 3,5-F_2_Y HPr. The structure of HPr (PDB id: 1QR5) is shown above the ^19^F spectra as cartoon representation, with Y6 side chain depicted as magenta sticks. b) On-resonance ^19^F *R*_1*ρ*_ dispersion profiles recorded at 16.4 T at the same temperatures as in a), with best fits (solid curves) shown as a function of the applied spin-lock field *ω*_SL_/2π. c) Eyring plot of *k*_flip_ derived from ^19^F *R*_1*ρ*_ RD data. The solid line is the least-squares fit, and the shaded region indicates the fit uncertainty. The errors of resulting activation parameters (*ΔH*^‡^ and *ΔS*^‡^) were estimated from Monte Carlo simulations. d) Experimental (black) and simulated (magenta)^19^F spectra for 2,6-F_2_Y6 in Y64F HPr from 35 °C to 55 °C, with *k*_flip_ values indicated for each temperature. e) The decay of auto (AA) and build-up curves of exchange (AB) peaks in ^19^F-^19^F EXSY experiment at varying mixing times for 2,6-F_2_Y6 in Y64F HPr at 10 °C. Solid lines are best fits for magnetization originating in the A and B states (AA and AB) to the equations in “Materials and Methods”. Peak volumes are normalized such that the peak volume at zero mixing time is 1.0. Inset shows the overlaid ^19^F-^19^F EXSY spectra recorded with a mixing time of 0 s (black) and 0.2 s (magenta). f) Eyring plot of *k*_flip_ rates for 2,6-F_2_Y6 in Y64F HPr from ^19^F LS (black) and EXSY (magenta), with resulting *ΔH*^‡^ and *ΔS*^‡^ shown in the figure and provided in Table 3.

In sharp contrast, the two ring-flipping states for 2,6-F_2_Y HPr are in slow exchange from 0 °C to 40 °C, rather than in fast exchange as observed for 3,5-F_2_Y HPr. The shift in exchange regime can be rationalized by larger *Δω*_AB_ values and/or slower *k*_flip_ rates for 2,6-F_2_Y6 compared to 3,5-F_2_Y6. Indeed, the *Δω*_AB_ for 2,6-F_2_Y6 at 10 °C is 2.3 ppm, significantly larger than that for 3,5-F_2_Y6 determined from *R*_1*ρ*_ RD (0.4 ppm). To quantitatively compare the *k*_flip_ rates, we performed ^19^F LS analysis using Y64F HPr mutant to alleviate spectral overlap (Figure 3d, Table 2) and confirmed that the Y64F mutation minimally affect Y6 ring flips (Table S3). Because *k*_flip_ can be underestimated in slow-exchange regime (*k*_flip_ ≪ ^19^F *R*_2,0_, *Δω*_AB_) due to its anticorrelation with ^19^F *R*_2,0_, as borne out by our Monte Carlo simulation studies (Figure S10), we performed ^19^F-^19^F EXSY experiments to accurately determine *k*_flip_ rates at low temperatures (Figure 3e). The joint use of *k*_flip_ rates from ^19^F LS analysis and EXSY enabled Eyring analysis over a broad temperature range (Figure 3f), yielding *ΔH*^‡^ = 145 ± 1 kJ·mol^-1^ and *ΔS*^‡^ = 277 ± 2 J·mol^-1^ K^-1^, both substantially higher than those determined for 3,5-F_2_Y6 (Table 3). Given that 2,6-F_2_Y incorporation slows Y6 ring flip by three magnitude orders, as observed for Y3 in GB1, we speculate that 2,6-F_2_Y can act as a general ring-flip “brake”, thereby facilitating detection of fast ring flips by placing them in a slower NMR timescale.

### Detection of Tyrosine ring flip in Ubiquitin by 3,5-F_2_Y and 2,6-F_2_Y

A previous study by Wand and coworkers concluded that the Y59 ring flip in Ub is absent below the transition temperature (30 °C) by ^13^C relaxation-based model-free analysis, which reports on the protein motions faster than *τ*_c_. Molecular simulations at 80 °C also failed to capture tyrosine ring flips in Ub.^[33]^ However, Akke and Weininger suggested an alternative interpretation of the ^13^C relaxation studies, namely that the Y59 ring flip occurs on a timescale significantly slower than *τ*_c_ below the transition temperature and thus escapes detection by model-free analysis.^[10]^ To test this hypothesis, we incorporated 2,6-/3,5-F_2_Y into Ub and found that neither labeling perturbs the protein structure, thermostability and backbone dynamics (Figures S11–12). The ^19^F peak of 3,5-F_2_Y59 in Ub at 5 °C (185 Hz) is significantly broader than those in GB1 and HPr, indicating the presence of ring-flip motion on the µs–ms timescale (Figure 4a). Therefore, we collected on-resonance ^19^F *R*_1*ρ*_ data and determined *k*_flip_ from 5 °C to 15 °C (Figure 4b, Table 1), supporting the hypothesis that the Y59 ring flip occurs on a timescale (*τ*_flip_ ≈ 15 µs) slower than *τ*_c_.

**Figure 4.**
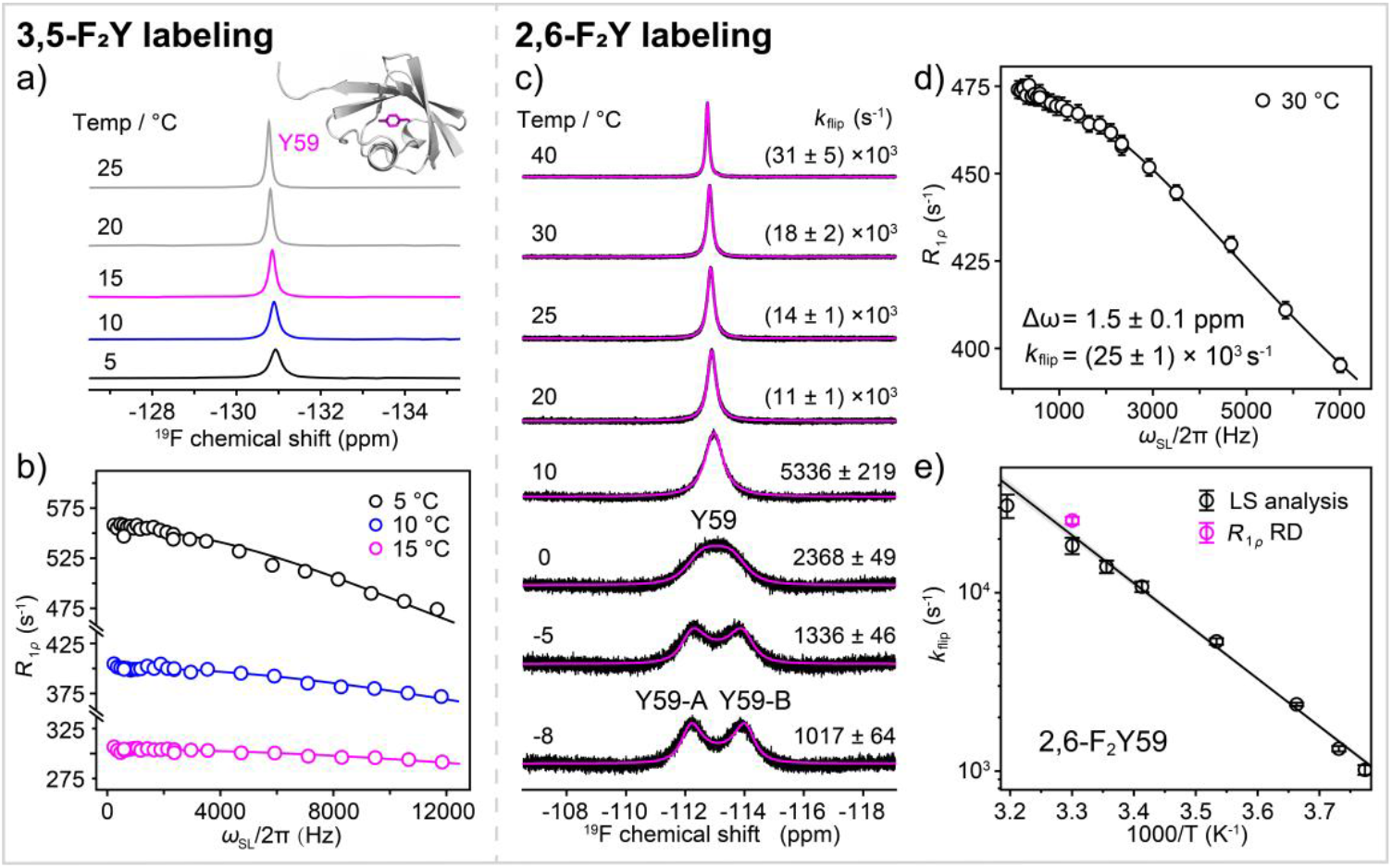
Ring-flip dynamics of Y59 in 3,5/2,6-F_2_Y Ub. a) Experimental ^19^F spectra for 3,5-F_2_Y59 in Ub recorded at 5 °C (black), 10 °C (blue), 15 °C (magenta), 20 °C (gray), and 25 °C (gray). The Ub structure (PDB id: 1UBQ) is shown above as a cartoon, with tyrosine side chains depicted as magenta sticks. b) On-resonance ^19^F *R*_1*ρ*_ dispersion profiles recorded for 3,5-F_2_Y59 at 16.4 T and at 5 °C (black), 10 °C (blue) and 15 °C (magenta). Solid curves are best fits of *R*_1*ρ*_ as a function of the applied spin-lock field *ω*_SL_/2π. c) Experimental (black) and simulated (magenta) ^19^F spectra for 2,6-F_2_Y59 in Ub at temperatures from -8 °C to 40 °C, with the corresponding *k*_flip_ indicated for each temperature. d) On-resonance ^19^F *R*_1*ρ*_ RD profile for 2,6-F_2_Y59 in Ub recorded at 30 °C, with the fitted chemical shift difference (1.5 ± 0.1 ppm) and *k*_flip_ ((25 ± 1) × 10^3^ s^-1^) indicated. e) Eyring plot of *k*_flip_ for 2,6-F_2_Y59 from ^19^F LS analysis. The solid line is a least-squares fit. The uncertainties of *k*_flip_ were estimated from Monte Carlo simulations. The *k*_flip_ from *R*_1*ρ*_ RD data at 30 °C is shown in magenta for comparison. The resulting activation parameters *ΔH*^‡^ and *ΔS*^‡^ values are given in Table 3.

To further confirm the existence of ring flips in Ub, we collected ^19^F spectra for 2,6-F_2_Y labeled Ub at temperatures from -8 to 40 °C (Figure 4c). The temperature-dependent ^19^F spectra for 2,6-F_2_Y Ub exhibit the characteristic NMR lineshapes of a two-state exchange with equal populations traversing from the slow- to fast-exchange regime, clearly indicating ring-flip motion. At -8 °C, we observed two distinct peaks of equal volume separated by 1.7 ppm (800 Hz), which coalesce around 0 °C. At the coalescence temperature, *k*_flip_ can be quantitatively determined to be 1777 s^-1^ 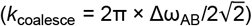.^[62-63]^ We then performed ^19^F LS analysis to determine *k*_flip_, fitting the chemical shift difference (*Δω*_AB_) at each temperature with a temperature coefficient (10 ± 1 ppb/°C) (Figure 4c), rather than assuming a common *Δω*_AB_. We validated the lineshape fitting with ^19^F *R*_1*ρ*_ RD data at 30 °C, and both the *R*_1*ρ*_-derived *Δω*_AB_ and *k*_flip_ are in excellent agreement with those derived from ^19^F LS analysis (Figure 4d, Table 2). In addition, *k*_flip_ at 0 °C was determined to be 2368 s^-1^, consistent with *k*_coalesce_ (1777 s^-1^). Interestingly, the activation enthalpy (*ΔH*^‡^ = 49 ± 1 kJ·mol^-1^) and entropy (*Δ*S^‡^ = -1 ± 4 J·mol^-1^ K^-1^) from Eyring analysis are significantly lower than those reported for other aromatic residues (Figure 4e), implying weaker stabilizing interactions and larger conformational freedom for Y59 in the ground state, in agreement with the Ub structure. The *k*_flip_ for 2,6-F_2_Y59 (5×10^3^ s^-1^ by LS analysis) is 20-fold slower than that for 3,5-F_2_Y59 ((90 ± 20) ×10^3^ s^-1^ by ^19^F *R*_1*ρ*_ RD) in Ub at 10 °C (Table 2). Unlike Y3 in GB1 and Y6 in HPr, Y59 is located in a loop region of Ub, where intra-residue steric repulsion may be mitigated by subtle backbone conformational rearrangements, resulting in a reduced hampering effect. Although 2,6-F_2_Y slows tyrosine ring flips in proteins, it nicely places the ring flips in slow-to-intermediate exchange regime, thereby facilitating detection of ring flips that would otherwise escape characterization due to large *k*_flip_ and/or small *Δω*_AB_.

### Determination of tyrosine ring-flip rates in living cells

Local unfolding has been suggested to facilitate aromatic ring-flip motions in globular proteins.^[12]^ In addition, previous studies have shown that cellular environments can alter protein folding stabilities by chemical interactions and macromolecular crowding effects.^[64-67]^ We therefore speculated that the ring-flip dynamics, reporting on the local conformational fluctuations, may differ in living cells compared to buffer. To test this hypothesis, we determined tyrosine ring-flip rates for two globular proteins, Y64F HPr and Y33F/Y45F GB1 (GB1^mut^), in *X. laevis* oocytes.

We used 2,6-F_2_Y labeling for in-cell ring-flip studies, because two well-resolved resonances are observed for Y64F HPr and GB1^mut^, and *k*_flip_ rates can be accurately determined by ^19^F-^19^F EXSY experiments in cells, whereas ^19^F relaxation dispersion experiments would be challenging due to the peak broadening.^[68]^ Notably, large ^19^F *R*_2,0_ and inhomogeneous peak broadening effects in *X. laevis* oocytes also complicate quantitative ^19^F LS analysis. 2,6-F_2_Y GB1^mut^ and 2,6-F_2_Y Y64F HPr were microinjected into *X. laevis* oocytes, and 1D in-cell ^19^F NMR spectra were recorded from 10 °C to 40 °C. At 40 °C, we observed significant peak broadening for 2,6-F_2_Y Y64F HPr, clearly indicating that tyrosine ring flips become faster with increasing temperature (Figure 5), whereas the ^19^F peak for Y37 sharpened as *τ*_c_ is decreased. The temperature-dependent ^19^F lineshapes in oocytes for 2,6-F_2_Y Y64F HPr closely resemble those in buffer. To quantitatively compare *k*_flip_ in cells and in buffer, we collected ^19^F-^19^F EXSY spectra for 2,6-F_2_Y Y64F HPr and GB1^mut^ at 10 °C and 25 °C in *X. laevis* oocytes and confirmed that the oocytes are viable and no protein leakage occurred during the in-cell experiments (Figure 5, Figure S13a–b). High-quality ^19^F-^19^F EXSY spectra were obtained at various mixing times, and *k*_flip_ rates were fitted with high accuracy. The *k*_flip_ values for 2,6-F_2_Y6 in Y64F HPr and 2,6-F_2_Y3 in GB1^mut^ in *X. laevis* oocytes are essentially the same as those measured in buffer (Table S3). Therefore, no significant change in ring-flip dynamics was found for two proteins in oocytes. Since macromolecular crowding stabilizes proteins while weak chemical interactions destabilize them, these opposing effects may cancel each other out, resulting in no net *k*_flip_ change. The *k*_flip_ may be affected more in human cells or *E. coli*, which are more crowded than *X. laevis* oocytes.

**Figure 5.**
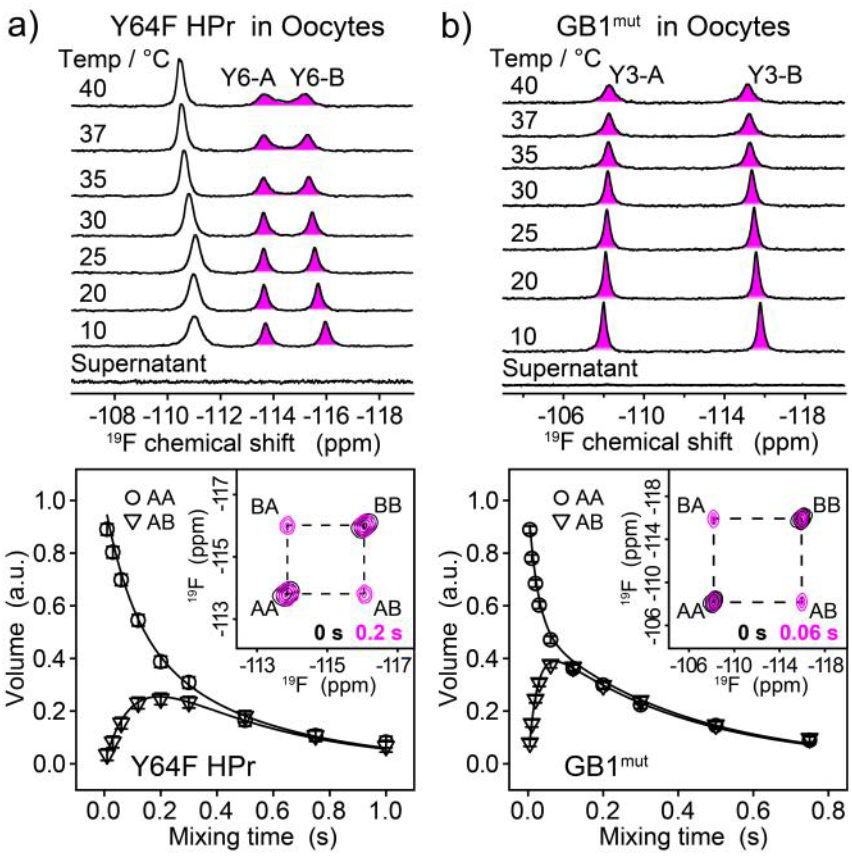
*k*_flip_ measurements in *X. laevis* oocytes. ^19^F spectra (top) and ^19^F-^19^F EXSY experiments (bottom) for a) 2,6-F_2_Y Y64F HPr and b) 2,6-F_2_Y GB1^mut^ in *X. laevis* oocytes. ^19^F spectra were recorded at temperatures ranging from 10 °C to 40 °C. Two ring-flipping states of 2,6-F_2_Y6 in Y64F HPr and 2,6-F_2_Y3 in GB1^mut^ are indicated as shaded areas in magenta. The ^19^F spectrum for the supernatant after in-cell experiments is shown at bottom to confirm the absence of protein leakage. Plots of the auto (AA) and exchange (AB) peak volumes as a function of mixing times at 10 °C are illustrated. The errors in peak volumes were estimated from spectrum noise. Solid lines are best fits of experimental data to the equations in “Materials and Methods”. Insets show the overlaid EXSY spectra with a mixing time of 0 s (black) and 0.2 s (magenta) for 2,6-F_2_Y6 in Y64F HPr, and 0 s (black) and 0.06 s (magenta) for 2,6-F_2_Y3 in GB1^mut^, with auto peaks of (AA and BB) and exchange peaks (AB and BA) indicated by dashed lines.

Synthetic polymers have been shown to stabilize proteins via macromolecular crowding effects. Indeed, ring flips in GB1^mut^ and Y64F HPr decreased in Ficoll 70K, possibly because of the increased protein stability (Figure S13c–d, Table S3). Together, we show that ring-flip dynamics persists and were not perturbed in *X. laevis* oocytes, but are slightly slowed in Ficoll 70K. More proteins and cell systems are needed to conclude the cellular effects on aromatic ring-flip dynamics in the future. Importantly, we show that the in-cell *k*_flip_ rates can be accurately measured by ^19^F-^19^F EXSY, which is impossible by ^1^H and ^13^C NMR due to non-specific interactions and cellular background.

## Conclusion

We have introduced two novel ring-flip ^19^F probes, 3,5-F_2_Y and 2,6-F_2_Y, which can be efficiently and cost-effectively incorporated into proteins for studying tyrosine ring-flip dynamics. The *k*_flip_ values for 3,5-F_2_Y in GB1 and HPr are on the same order as those reported in previous ^1^H and ^13^C studies,^[11, 13]^ and the observed discrepancies likely arise from differences in experimental conditions, the NMR techniques employed, and fluorination effects. Aromatic ^19^F atoms are poor hydrogen-bond acceptors, and previous work has shown that fluorinated aromatic amino acids minimally impact protein structures.^[69-72]^ However, fluorine’s strong electron-withdrawing effects can alter the charge distribution of the aromatic ring and thereby perturb aromatic π-π interactions, particularly for tyrosines in aromatic clusters. This is supported by a similar *k*_flip_ for isolated 3,5-F_2_Y6 in HPr and 5–8 times faster *k*_flip_ for 3,5-F_2_Y3 and 3,5-F_2_Y45 in the GB1 aromatic cluster. Using 3,5-F_2_Y labeling, we captured and quantitatively characterized Y59 ring flips in ubiquitin (τ_flip_ ≈ 15 µs), which had been previously suggested to be absent below transition temperature (30 °C). Remarkably, we observed a dramatic slowdown of tyrosine ring flips in three different 2,6-F_2_Y labeled proteins, indicating that 2,6-F_2_Y generally hampers the ring flips, likely through the steric repulsion between ^19^F atoms and backbone atoms. To our knowledge, the modulation of ring flips without extreme conditions (*e*.*g*., high pressure and low temperatures) has only been reported in a peptide ssNMR study, in which the flip of a phenylalanine ring was hindered by an adjacent methyl group.^[73]^ Moreover, we demonstrate the utility of 2,6-F_2_Y labeling to measure *k*_flip_ in living *X. laevis* oocytes for the first time, which is almost impossible for ^1^H or ^13^C labeling.

Taken together, these results establish 3,5-F_2_Y as a generally native-like ring-flip probe for characterizing ring flip dynamics, with only moderate perturbations in tightly packed aromatic clusters, whereas 2,6-F_2_Y serves as a novel ring-flip “brake” that effectively “knocks down” the ring-flip dynamics without perturbing protein folding. Given that ring-flip motion has long been recognized as a hallmark of protein “breathing” motion and was until recently implicated in mediating protein “breathing” motion and receptor activation, 2,6-F_2_Y provides a unique tool to directly modulate and dissect the effects of ring flips on protein structure, dynamics, and interactions. Furthermore, the absence of ^19^F in cells renders 3,5-F_2_Y and 2,6-F_2_Y well suited for quantifying ring-flip dynamics in living cells and elucidating cellular environmental effects on protein “breathing” motions. Our studies therefore open a new avenue for probing and modulating aromatic ring-flip dynamics both *in vitro* and in cells.

## Acknowledgements

This work was supported by the Chinese Academy of Sciences (XDB0540000 and YSBR-068), the National Natural Science Foundation of China [grants 22374156, 21925406], and the Ministry of Science and Technology of China [grants 2021YFA1302600].

