## supplemental information for "Characterizing Tyrosine Ring Flips in Proteins by ^19^F NMR"

### Experimental Procedures

#### Enzymatic synthesis of 3,5-/2,6-$F_2Y$

2,6-difluorophenol, 3,5-difluorophenol, pyridoxal-5′-phosphate (PLP), Celite® 545 were purchased from Aladdin (Shanghai, China). $^{15}NH_4Cl$ was purchased from Cambridge Isotope Laboratories, Inc. (Andover, USA). Sodium pyruvate and ethyl acetate were purchased from Sinopharm Chemical Reagent Co., Ltd. (Shanghai, China).

The enzyme tyrosine phenol lyase (TPL) was obtained as previously described with slight modifications.[1] The gene encoding TPL from C. *freundii* (NCBI accession No. L10821.1) was synthesized by Sangon Biotechnology Co., Ltd. (Shanghai, China) and cloned into the pET-28a vector with restriction sites BamHI/XhoI under control of the T7 promoter. TPL purification was performed using HisTrap Ni-NTA column at 4 °C using Ni A buffer (50 mM $KH_2PO_4$-$K_2HPO_4$, 500 mM NaCl, 10 mM imidazole, pH 7.5) and Ni B buffer (50 mM $KH_2PO_4$-$K_2HPO_4$, 500 mM NaCl, 500 mM imidazole, pH 7.5). The purified TPL was buffer exchanged to 20 mM $KH_2PO_4$-$K_2HPO_4$, pH 7,5, and stored at -80 °C.

3,5-/2,6-$F_2Y$ were synthesized as described.[2] The final products were identified by $^{19}F$ NMR and ESI-MS. The $F_2Y$ yield ($Y_{F_2Y}$) was calculated as below:

$$Y_{F_2Y} = \frac{Y_{F_2Y,T}\ \alpha\ I_{F_2Y,2}\ V_2}{I_{F_2Y,1}\ V_1} \qquad \text{eq 1}$$

where $I_{F_2Y,1}$ and $I_{F_2Y,2}$ are the $^{19}F$ peak integrals of $F_2Y$, using 4-F phenylalanine as a reference, in the reaction mixture and in the final product, respectively. $V_1$ and $V_2$ are the volumes of the reaction mixture and final product, respectively. $\alpha$ is the reaction conversion rate, and $Y_{F_2Y,T}$ is the theoretical yield of $F_2Y$.

#### Protein expression and purification

The genes encoding GB1, Ub and HPr (*Staphylococcus carnosus,* UniProtKB P23534) were cloned into the pET-28a vector with restriction sites NcoI/XhoI under the T7 promoter. Mutations were introduced by site-directed mutagenesis. All primers were synthesized by Sangon Biotechnology Co., Ltd. (Shanghai, China). The plasmids encoding the above proteins were transformed into *E. coli* BL21(DE3). $^{19}F$ labeled GB1 and its mutants (Y3F, Y33F, Y45F, Y33F/Y45F) and Ub were expressed and purified as described for 3FY incorporation.[3-4] HPr and its mutants (Y6F, Y37F, Y64F, Y37F/Y64F) were purified as previously described with slight modifications.[5] Briefly, the cell pellets were resuspended in 50 ml buffer A (20 mM sodium phosphate, pH 7.0) containing 500 µL protease inhibitor cocktail (100 x) and 1mg DNase. Cells were lysed by 3 passes through an ATS high-pressure homogenizer at 900 bar. The cell debris was removed by centrifugation at 20,000 rpm for 30 min at 4 °C. The supernatant was subjected to a DEAE Fast Flow column (GE Healthcare) and eluted with a linear gradient consisting of buffer A and buffer B (20 mM sodium phosphate, 1 M NaCl, pH 7.0), followed by size exclusion chromatography pre-equilibrated with buffer C (20 mM Tris, 150 M NaCl, pH 7.5). All proteins were desalted, lyophilized, and stored at -80 °C.

**NMR sample preparation**

For *in vitro* NMR, the measurements below 0 °C were achieved using eight 1.1 mm o.d. glass capillary tubes inserted into a regular 5 mm NMR tube.[6-7] Samples were centrifuged at 16,000 g for at least 30 minutes before loading into capillary tubes to remove all potential freezing-nucleation sites from the aqueous solutions. 2,6-/3,5-$F_2$Y GB1 (2-3 mM) were prepared in 20 mM MES (pH 6.5) with 100% (v/v) $D_2O$. 2,6-$F_2$Y Ub (7.3 mM) was prepared in 20 mM HEPES (pH 7.5) with 100% (v/v) $D_2O$. $^{19}$F signal assignments were achieved with 0.3 mM protein samples. For $^{15}$N relaxation and $^{19}$F-$^{19}$F EXSY measurements, 1 mM proteins were prepared in 20 mM MES (pH 6.5), 100 mM KCl, and 10% (v/v) $D_2O$ for GB1, 20 mM HEPES (pH 7.5), 150 mM NaCl, 10% (v/v) $D_2O$ for HPr and Ub, respectively, using regular 5 mm NMR tubes. Samples for $^{19}$F $R_{1\rho}$ experiments (1 mM) were prepared in same buffer without salt.

For in-cell NMR, *X. laevis* oocytes were prepared as described,[8] and 10 mM 2,6-$F_2$Y labeled protein was used for microinjection to produce approximately 0.4 mM protein in *X. laevis* oocytes for in-cell experiments. The protein samples for studying macromolecule crowding effects were prepared by adding 300 g/L Ficoll 70K to the dilute samples.

**NMR spectroscopy**

NMR spectra were recorded on Bruker Avance spectrometers at 500 MHz equipped with a double-resonance broadband probe or at 600, 700, and 850 MHz equipped with triple-resonance cryogenic probes. $^{19}$F temperature-variable spectra from -10 °C to 70 °C, were recorded at 500 MHz with 256–512 scans. Notably, capillary samples were cooled slowly (-1 °C every 3 minutes) to reach temperatures well below 0 °C and maintain a supercooled state without freezing the sample. $^{19}$F temperature-variable spectra from 5 °C to 40 °C and $^{19}$F NMR spectra for signal assignment were recorded at 600 MHz with 256-1024 scans.

$^{19}$F $R_2$ rates were measured by CPMG at 0 and 10 °C for Y3 of 2,6-$F_2$Y Y33F/Y45F GB1, and at -10 °C, 0 °C and 10 °C for Y6 of 2,6-$F_2$Y Y37F/Y64F HPr. $R_2$ relaxation rates were obtained by fitting the intensity decay to a single-exponential functions ($I_t = I_0 \times \exp(-R_2 \times t)$).[9]

$^{19}$F $R_{1\rho}$ relaxation dispersion (RD) spectra were recorded at 700 MHz equipped with a QCI $^{19}$F probe using the published pulse sequence[10]. $^{19}$F $R_{1\rho}$ RD profiles were recorded for Y3 of 3,5-$F_2$Y GB1 at 5.0, 7.5, 10.0 and 12.5 °C; for Y45 of 3,5-$F_2$Y GB1 at 20.0, 25.0 and 30.0 °C; for Y6 of 3,5-$F_2$Y HPr at 5.0, 7.5, 10.0, 12.5 and 15.0 °C; for Y59 of 3,5-$F_2$Y Ub at 5, 10 and 15 °C; and for Y59 of 2,6-$F_2$Y Ub at 30 °C using a relaxation delay of 1.5 s and 6-8 spin-lock times $T_{SL}$. For Y45 of 3,5-$F_2$Y GB1 and Y59 of 3,5-$F_2$Y Ub, 25 spin-lock field strengths ($\omega_{SL}$) between 100 and10000 Hz were used, whereas $\omega_{SL}$ between 100 and 6000 Hz were used for all other samples. The 90° pulse width and the spin lock fields were calibrated for each experiment with 2 mM trifluoroacetic acid (TFA). Importantly, the $^{19}$F carrier frequency was set at the center of the peak of interest in the $^{19}$F spectrum for each protein. For Y3 in 3,5-$F_2$Y GB1 and Y6 in 3,5-$F_2$Y HPr, $^{19}$F $R_{1\rho}$ RD data were recorded at two magnetic fields (14.1 and 16.4 T) for robust fitting of $k_{ex}$ and $\Delta\omega_{F,AB}$.

$^{19}$F-$^{19}$F EXSY spectra were recorded at 600 MHz at 10 °C and 25 °C with 16 scans for dilute samples and 48–64 scans for *X. laevis* oocytes, a relaxation delay of 1.5 s, and 2048 × 32 ($t_2 \times t_1$) complex points. The mixing times for 2,6-$F_2$Y GB1 and Y33F/Y45F GB1 were 0, 5, 10, 20, 30, 60, 120, 200, 300, 500 and 750 ms at 10 °C and 0, 0.5, 2, 5, 10, 30, 60, 120, 200, 300 and 500 ms at 20–35 °C. For 2,6-$F_2$Y HPr and Y37F/Y64F HPr, the mixing times were 0, 10, 30, 60, 120, 200, 300, 500, 750 and 1000 ms at 10 °C and 0, 2, 5, 10, 20, 30, 60, 120, 200, 300 and 500 ms at 25 °C.

$^1$H-$^{15}$N HSQC spectra were recorded at 600 or 700 MHz at 25 °C with 8 scans, a 2 s relaxation delay, and 2048 × 128 ($t_2 \times t_1$) complex points. $^1$H-$^{15}$N SOFAST HMQC spectra for *X. laevis* oocytes were recorded at 850 MHz at 25 °C with 128–512 scans and 2048 × 256 ($t_2 \times t_1$) complex points. $^{15}$N $T_1$/$T_2$ relaxation experiments were recorded at 600 MHz at 25 °C. For $T_1$ measurements, relaxation delays were 0.01, 0.1, 0.2, 0.35, 0.5, 0.7, 1.0, 1.5, 2.0 and 2.5 s. For $T_2$ measurements, the relaxation delays were 0.017, 0.051, 0.085, 0.153, 0.220, 0.322, 0.424, 0.543 and 0.678 s. NMR spectra were processed with TopSpin 3.6.5 and analyzed with NMRPipe[11].

**Mass Spectrometry (MS)**

Spectra were recorded on an Agilent Technologies 6530 (Q-TOF) with an ESI source connected to an Agilent 1200 series HPLC equipped with a ZORBAX SB-C3 column (5μm, 4.6×150 mm). Approximately 20 μM of sample was injected, proteins were eluted with a 1 mL/min linear gradient from 5% to 40% solvent B (acetonitrile) in 15 minutes (solvent A: 0.1% (v/v) formic acid in water). The ESI source was operated in positive-ion mode. Data were processed with Agilent Masshunter Qualitative Analysis.

**Circular Dichroism (CD) spectroscopy**

CD spectra and Melting curves were recorded on a Chirascan spectropolarimeter (Applied Photophysics Limited. UK) equipped with a Quantum Northwest TC125 temperature controller, using 15 μM protein samples. CD spectra were acquired at the wavelength from 190 to 260 nm at 25 and 80 °C with a step size of 1 nm, a time per point of 0.5 s, and a bandwidth of 1 nm. Melting curves for HPr, 3,5-$F_2$Y HPr and 2,6-$F_2$Y HPr were measured at 208 nm with a temperature ramp of 0.5 °C/min.

**$^{19}$F lineshape analysis**

The $^{19}$F lineshape analysis for quantifying ring flips was performed by numerically solving the Bloch-McConnell equation for two-state exchange in an uncoupled spin system described as follow:

$$\frac{\mathrm{d}}{\mathrm{d}t}\begin{pmatrix} M_A \\ M_B \end{pmatrix} = -(\mathrm{i}\mathbf{L} + \mathbf{R} + \mathbf{K})\begin{pmatrix} M_A \\ M_B \end{pmatrix} \qquad \text{eq 2}$$

$$\mathbf{L} = \begin{bmatrix} \Omega_A & 0 \\ 0 & \Omega_B \end{bmatrix};\ \mathbf{R} = \begin{bmatrix} -R_{2A} & 0 \\ 0 & -R_{2B} \end{bmatrix};\ \mathbf{K} = \begin{bmatrix} -k_{AB} & k_{BA} \\ k_{AB} & -k_{BA} \end{bmatrix};$$

Where $M_A$ and $M_B$ are the magnetizations of A or B state; $\Omega_{A,B}$ and $R_{2A,B}$ are chemical shifts and transverse relaxation rates of state A and B; and $k_{AB}$ $(= k_{BA})$ is the $k_{\mathrm{flip}}$ rate. **L**, **R** and **K** denote the Liouvillian, relaxation and exchange matrix, respectively. Diagonalization of the total matrix (i**L** + **R** + **K**) yields eigenvalues and eigenvectors, which are used to calculate the frequency domain spectrum following the method described previously[12-14]. The calculated spectrum is fitted to the experimental spectrum using least-squares method, with fixed $M_{A,0}$ = $M_{B,0}$ and $R_{2A}$ = $R_{2B}$; The population averaged chemical shift $\Omega_{ave}$ $(=\frac{\Omega_A+\Omega_B}{2})$ was obtained directly from the spectrum at each temperature and fixed during the fitting. Floating the chemical shift differences between A and B state $\Delta\Omega$ $(\Omega_A - \Omega_B)$ frequently produces the $\Delta\Omega$ values to the setting bounds, indicating the poor convergence. To address this, we instead optimized and varied $\Delta\Omega$ as a linear function of the temperatures using a $^{19}$F chemical shift temperature coefficient (ppb/k) during the fitting procedure. The goodness of the fit was indicated by reduced $\chi^2$, defined as $\chi^2/(N\text{-}K)$, where $\chi^2$ is computed as $\sum_i^N(\frac{I_i^{calc}-I_i^{exp}}{\sigma_i^{exp}})^2$, $N$ and $K$ denote the number of data points and the fitted parameters, respectively. The uncertainties of intensities were estimated from the standard deviation of the baseline region in the experimental $^{19}$F spectra.

According to our Monte Carlo simulations (Figure S10), $R_{2,A}$ and $\Delta\Omega_{AB}$ significantly affects the accuracy of the fitted $k_{flip}$ rates in the slow and fast exchange regimes, respectively. Accordingly, we performed $^{19}$F $R_{1\rho}$ RD experiments to independently validate the chemical shift difference $(\Delta\Omega_{AB})$ and $k_{flip}$ rates in the intermediate-to-fast exchange when $\Delta\Omega_{AB}$ can't be reliably determined. To account for $^{19}$F intrinsic $R_{2,0}$, we determined $R_{2,0}$ of the GB1 and HPr at low temperatures, where $R_{ex}$ contribution from slow chemical exchange can be completely suppressed by CPMG pulses with 2 kHz frequency (Table S1). $R_{2,0}$ for Ub is determined from $^{19}$F $R_{1\rho}$ RD experiment at 30 °C. Subsequently, $R_{2,0}$ for high temperatures were extrapolated from those measured at low temperatures based on the rotational correlation time ($\tau_c$) calculated by *HydroPro* program[15] since $^{19}$F $R_{2,0}$ is known to be dominated by chemical shift anisotropy, *i.e.*, proportional to the rotational correlation time, assuming the internal motions are unaffected by the temperatures. Finally, the uncertainties of $k_{flip}$ rates were estimated from 500 Monte Carlo runs by varying $R_{2,0}$ by ± 25%.

**On-resonance $^{19}$F $R_{1\rho}$ relaxation dispersion**

1D $^{19}$F $R_{1\rho}$ relaxation dispersion data were processed in MestReNova[16] and analyzed with in-house Python scripts. $^{19}$F $R_{1\rho}$ rates at each spin lock field were extracted by fitting peak intensities, measured at different $T_{SL}$ times, to a single-exponential decay. The resulting on-resonance $R_{1\rho}$ RD data were then fitted by numerically solving the Bloch-McConnell equations for describing chemical exchange in the presence of radio-frequency fields[17-18]. The populations were fixed at $p_A$ = $p_B$ = 0.5, while $k_{ex}$ (2 × $k_{flip}$) and $\Delta\omega_{F,AB}$ were optimized during the fitting through the least-squares optimization. The reduced $\chi^2$ was computed to assess the goodness of the fit. The uncertainties in the fitted parameters ($k_{ex}$ and $\Delta\omega_{F,AB}$) were estimated from the covariance matrix.

**Eyring analysis**

Activation parameters of the ring flips were determined using the Eyring equation through least-squares minimization of the linear fit of $log\ (\frac{k_{flip}}{T})$ versus $1/T$:[6, 19-20]

$$log\left(\frac{k_{\mathrm{flip}}}{T}\right) = -\frac{\Delta H^{\ddagger}}{2.303R} \times \frac{1}{T} + \frac{\Delta S^{\ddagger}}{2.303R} + log\left(\frac{k_{\mathrm{B}}}{h}\right) \qquad \text{eq 3}$$

where $k_B$, $h$, and $R$ are Boltzmann's constant, Planck's constant, and the universal gas constant, respectively; $T$ is the absolute temperature in Kelvin. $\Delta H^{\ddagger}$ and $\Delta S^{\ddagger}$ are the activation enthalpy and activation entropy, respectively, assumed to be independent of $T$. The uncertainties in the fitted $\Delta H^{\ddagger}$ and $\Delta S^{\ddagger}$ were estimated from 1000 Monte Carlo simulations.

**$^{19}$F-$^{19}$F EXSY analysis**

$^{19}$F-$^{19}$F EXSY experiments were recorded at various mixing times to quantify the exchange rate constants between A and B states. The auto and exchange peak volumes of 2,6-$F_2$Y residue ($I_{AA}$, $I_{BB}$, $I_{AB}$, $I_{BA}$) were extracted and fitted simultaneously using a least-squares procedure to extract the chemical exchange and $^{19}$F longitudinal relaxation rates as described previously:

$$I_{AA}(T) / I_{AA}(0) = [-(\lambda_2 - a_{11}) \exp(-\lambda_1 T) + (\lambda_1 - a_{11}) \exp(-\lambda_2 T)] / (\lambda_1 - \lambda_2) \qquad \text{eq 4}$$

$$I_{BB}(T) / I_{BB}(0) = [-(\lambda_2 - a_{22}) \exp(-\lambda_1 T) + (\lambda_1 - a_{22}) \exp(-\lambda_2 T)] / (\lambda_1 - \lambda_2) \qquad \text{eq 5}$$

$$I_{AB}(T) / I_{AA}(0) = [a_{21} \exp(-\lambda_1 T) - a_{21} \exp(-\lambda_2 T)] / (\lambda_1 - \lambda_2) \qquad \text{eq 6}$$

$$I_{BA}(T) / I_{BB}(0) = [a_{12} \exp(-\lambda_1 T) - a_{12} \exp(-\lambda_2 T)] / (\lambda_1 - \lambda_2) \qquad \text{eq 7}$$

where, $a_{11} = R_{1,A} + k_{AB}$, $a_{12} = -k_{BA}$, $a_{21} = -k_{AB}$, $a_{22} = R_{1,B} + k_{BA}$. $R_{1,A}$ and $R_{1,B}$ are the $^{19}$F longitudinal relaxation rates of the A and B states, respectively; $k_{AB}$ and $k_{BA}$ are the A to B and B to A flipping rate constants, respectively. $I_{AA}(T)$ and $I_{BB}(T)$ are peak volumes of the A and B auto peaks, and $I_{AB}(T)$ and $I_{BA}(T)$ are peak volumes of the exchange cross peaks arising from A to B and B to A at various mixing times T, respectively. $I_{AA}(0)$ and $I_{BB}(0)$ are the $^{19}$F magnetization for A and B states at zero mixing time. Because the populations of the A and B states are equal (0.5) for the 180° aromatic ring flips, the fitting is conducted under the constraint $k_{AB} = k_{BA} = k_{flip}$.

**Molecular dynamics simulations**

To obtain the free energy landscape of tyrosine ring flipping in the short peptide AYA, 1000 ns of Adaptively Biased Molecular Dynamics (ABMD) [21] simulations were performed using Amber 22 [22] for the wild-type AYA, as well as for its 2,6-/3,5-$F_2$Y labeled variants. The FF19SB force field [23] was applied to the peptide, while the General Amber Force Field (GAFF) [24] was used for the $^{19}$F-labeled tyrosine residues. The initial AYA structure was built in PyMOL.[25] The 3,5-$F_2$Y was modeled by replacing the HD1 and HD2 atoms with fluorine atoms, and similarly, the 3,5-$F_2$Y was created by replacing the HE1 and HE2 atoms. The peptide was then solvated in a TIP3P water box with a 15 Å buffer distance from the box edges. The system was neutralized with appropriate ions, and periodic boundary conditions were applied throughout the simulation. An initial minimization of 20,000 steps was carried out to relax the solvent with the peptide restrained. The system was heated to 298 K using Langevin dynamics, and pressure was maintained with a Monte Carlo barostat. Bonds involving hydrogen atoms were constrained using the SHAKE algorithm.[26] The Particle Mesh Ewald (PME) [27] method treated electrostatic interactions, and a 10 Å cutoff was applied for non-bonded interactions. Following equilibration, 1000 ns ABMD production runs were conducted for all three systems. The $\chi_2$ dihedral angle of tyrosine was chosen as the collective variable. The free energy profile as a function of the chi2 angle was subsequently generated using nfe-umbrella-slice in AmberTools.[28]

### Supplementary Figures and Tables

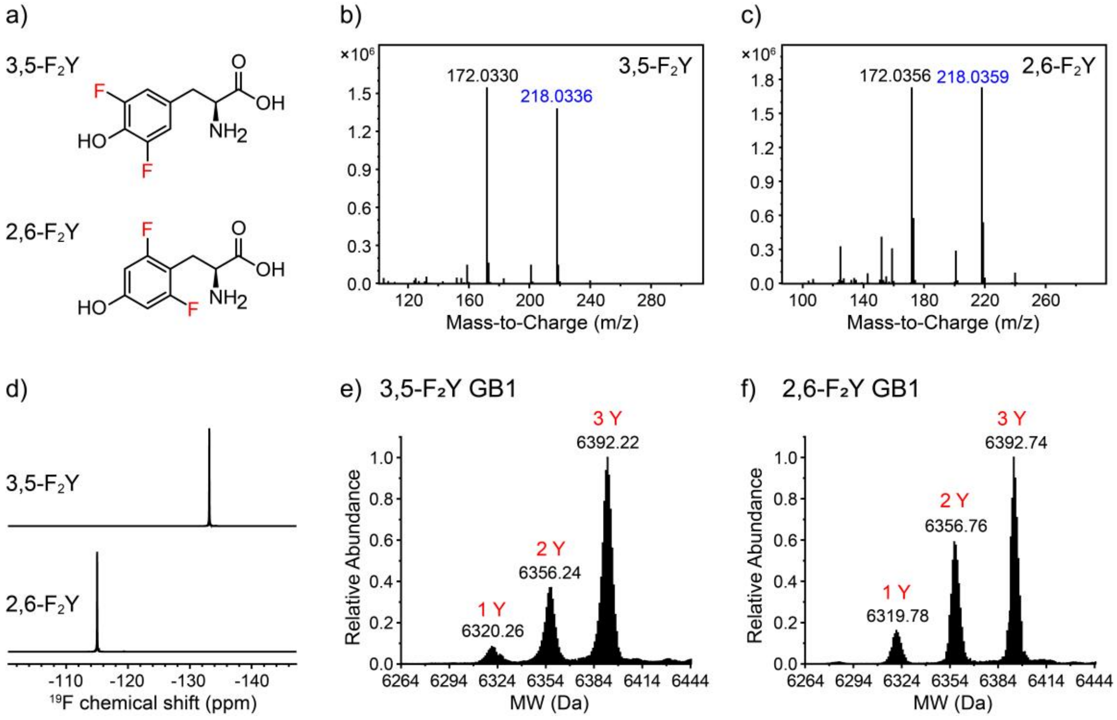

**Figure S1.** 3,5-/2,6-$F_2Y$ labeling of GB1. a) Chemical structures of symmetrical di-fluorotyrosines. (b–c) ESI-MS spectra of b) 3,5-$F_2Y$ and c) 2,6-$F_2Y$. The $[M+1]^+$ signals of 3,5-/2,6-$F_2Y$ are marked in blue. d) $^{19}F$ spectra of 3,5-$F_2Y$ (top) and 2,6-$F_2Y$ (bottom). (e–f) Mass spectra of e) 3,5-$F_2Y$ GB1 and f) 2,6-$F_2Y$ GB1. The red marks indicate the number of tyrosine residues successfully labeled with $F_2Y$. The yield of GB1 with fluorine substitution at all three sites are 63.0% and 53.2%, respectively.

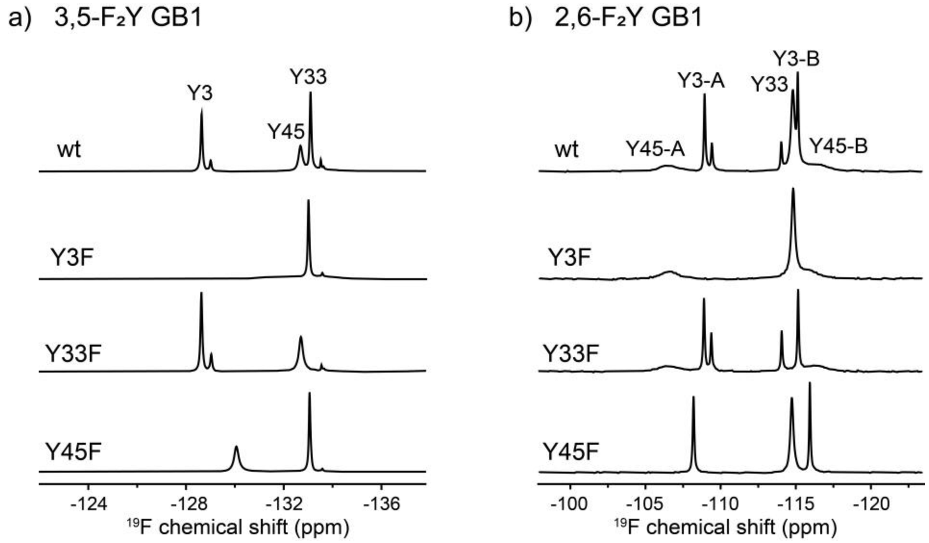

**Figure S2.** $^{19}F$ peak assignments for 3,5-/2,6-$F_2$Y GB1. (a–b) $^{19}F$ spectra of a) 3,5-$F_2$Y GB1 and b) 2,6-$F_2$Y GB1 and their Y to F variants (Y3F, Y33F, and Y45F).

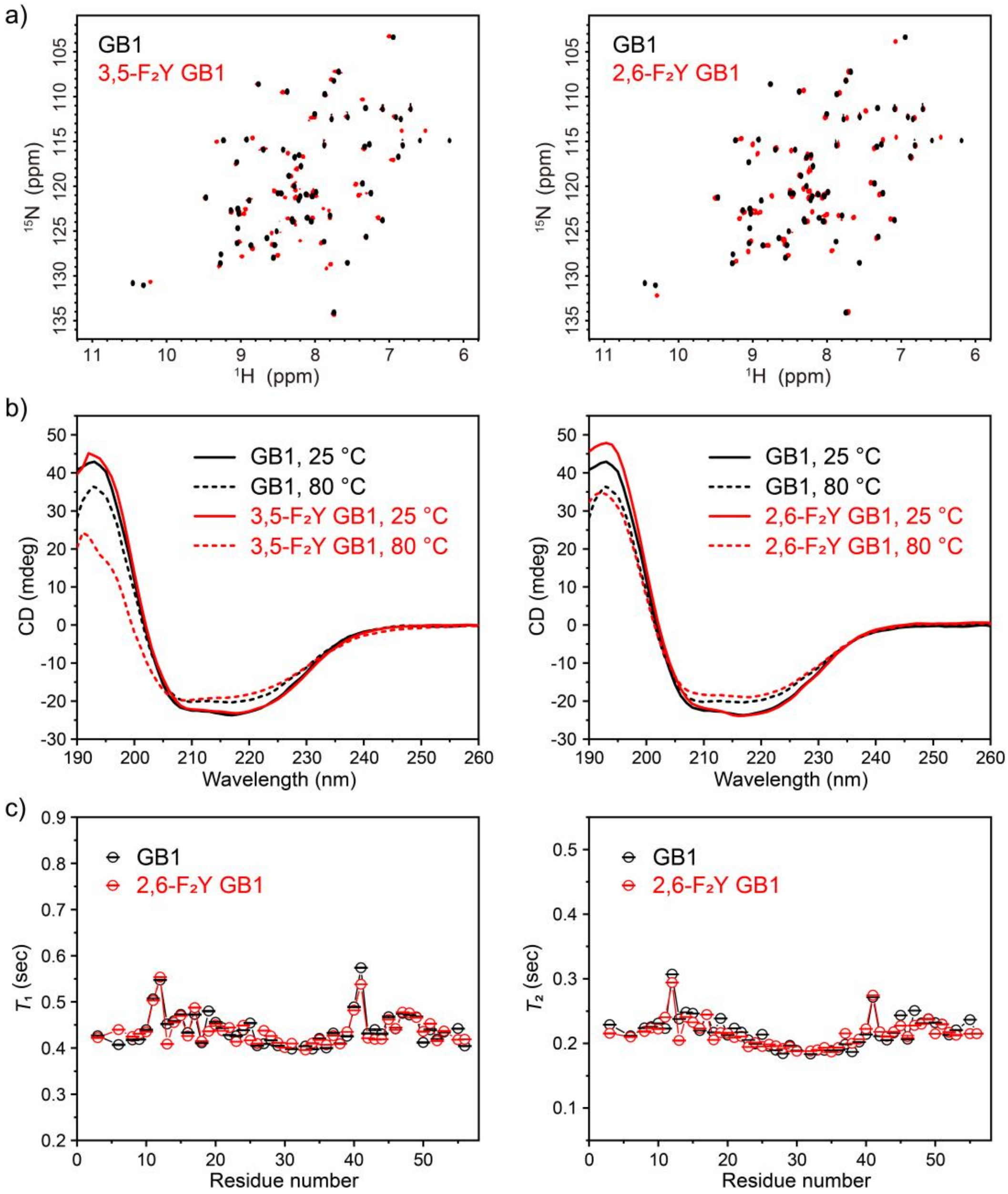

**Figure S3.** $^{1}$H-$^{15}$N HSQC spectra, CD spectra and $^{15}$N relaxation data for 3,5-/2,6-$F_2$Y GB1. a) Overlaid $^{1}$H-$^{15}$N HSQC spectra of GB1 with 3,5-$F_2$Y GB1 (left) and with 2,6-$F_2$Y GB1 (right). All spectra were recorded at 600 MHz at 25 °C with identical acquisition parameters. b) Overlaid CD spectra of GB1 (black) and 3,5-$F_2$Y GB1 or 2,6-$F_2$Y GB1 (red) at 25 °C (solid) and 80 °C (dashed), respectively. c) $^{15}$N $T_1$ and $T_2$ measurements for GB1 (black) and 2,6-$F_2$Y GB1 (red) at 600 MHz.

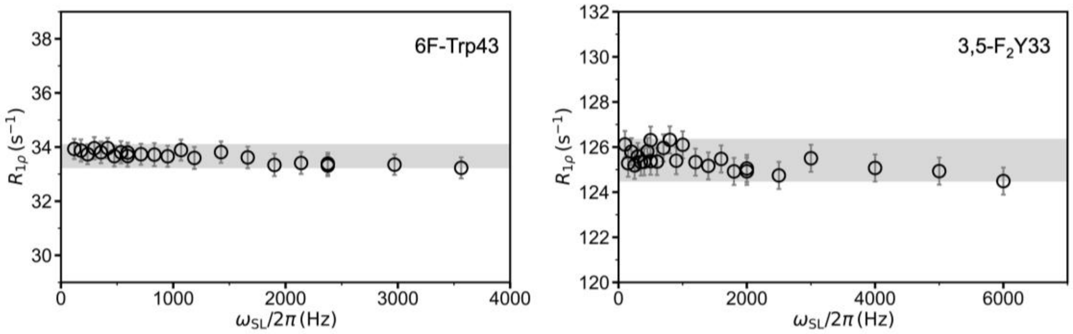

**Figure S4.** The [19]F $R_{1\rho}$ relaxation dispersion data for the aromatic side chain of Trp43 in 6FW GB1 (left), in which ring-flip motion is known to be absent, and Y33 in 3,5-$F_2$Y GB1 (right), which undergoes fast ring flips. The regions shaded in gray indicate the range corresponding to 2 × rmsd of the experimental $R_{1\rho}$ values.

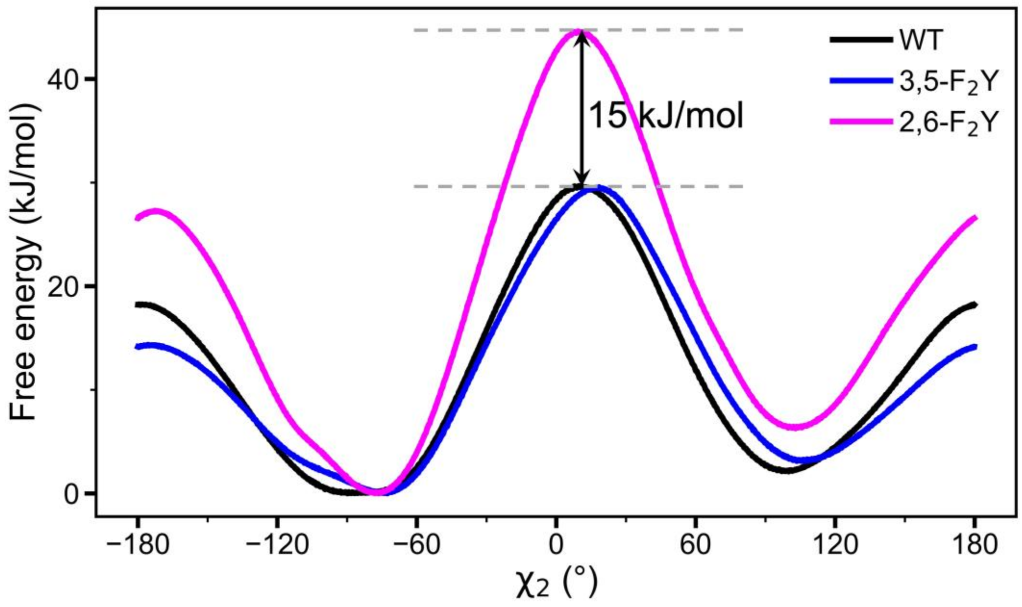

**Figure S5.** Free-energy profiles as a function of the $\chi_2$ dihedral angles for native (black) and fluorinated AYA tripeptides, 3,5-$F_2$Y (blue) and 2,6-$F_2$Y (magenta), during tyrosine ring flips, obtained from MD simulations. The transition-state free-energy difference ($\Delta\Delta G^{‡}$) for ring flips between 2,6-$F_2$Y and native Tyr (3,5-$F_2$Y) is indicated.

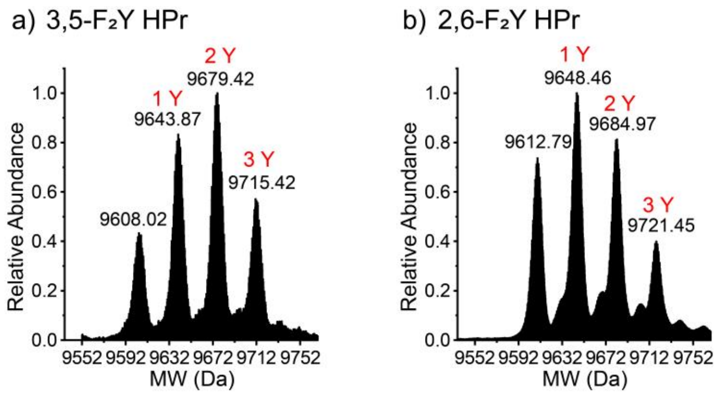

**Figure S6.** (a–b) Mass spectra of a) 3,5-$F_2$Y HPr and b) 2,6-$F_2$Y HPr. The red marks indicate the number of tyrosine residues successfully labeled with di-fluorotyrosine. The yield of proteins with fluorine substitution at all three tyrosine sites are 20.7% and 15.5%, respectively. An extra set of peaks are the adduct ions of $[M+Na]^+$.

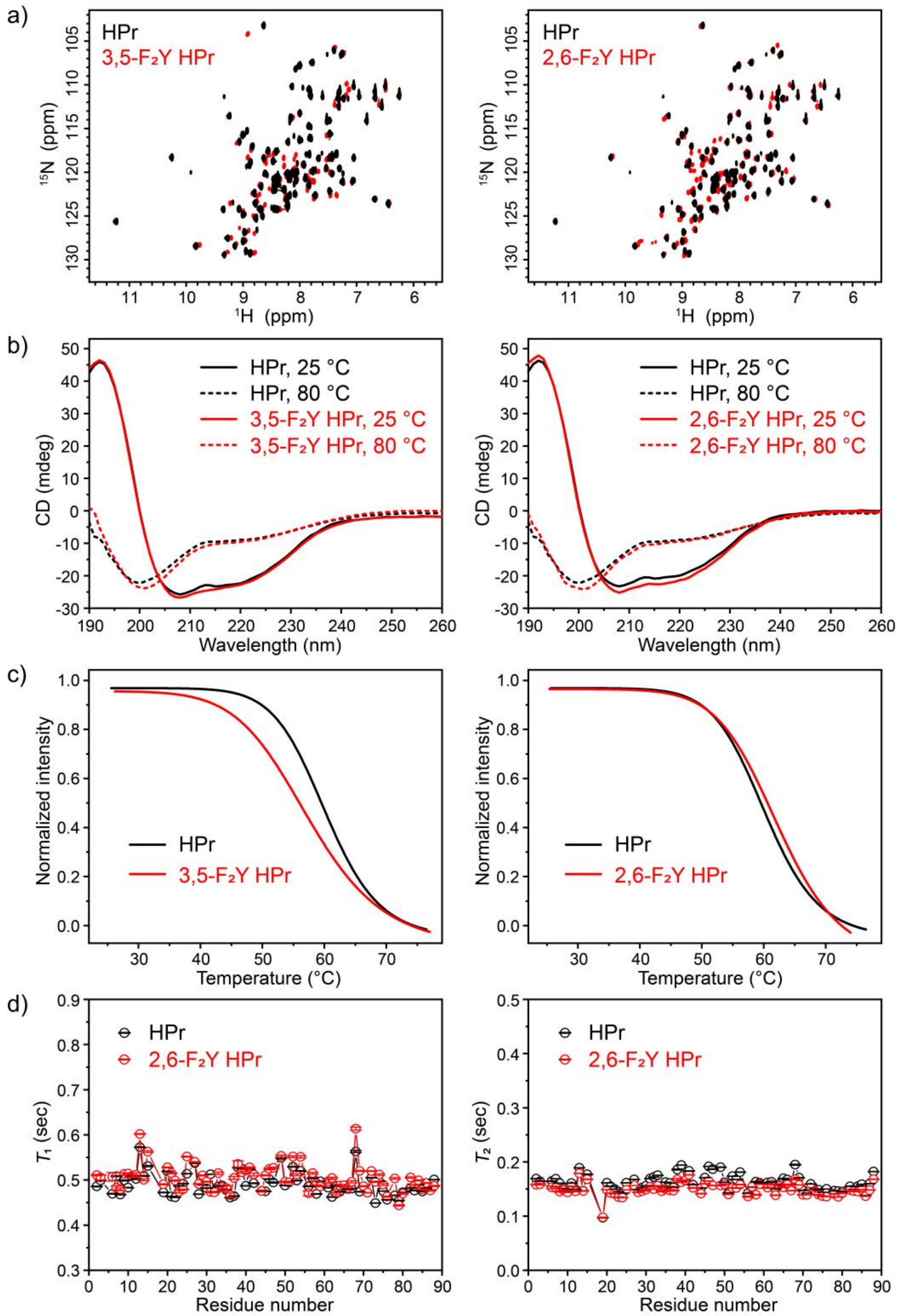

**Figure S7.** $^{1}$H-$^{15}$N HSQC spectra, CD spectra and $^{15}$N relaxation data for $F_2$Y HPr. a) Overlaid $^{1}$H-$^{15}$N HSQC spectra of HPr (black) and 3,5-$F_2$Y HPr or 2,6-$F_2$Y HPr (red). b) Overlaid CD spectra of HPr (black) and 3,5-$F_2$Y HPr or 2,6-$F_2$Y HPr (red) at 25 °C (solid) and 80 °C (dashed), respectively. c) The melting curves of HPr (black) and 3,5-$F_2$Y HPr or 2,6-$F_2$Y HPr (red). The $T_m$ values for HPr, 3,5-$F_2$Y and 2,6-$F_2$Y HPr are 60.2 ± 0.3, 57.5 ± 0.5, and 62.3 ± 0.5 °C, respectively. d) $^{15}$N $T_1$ and $T_2$ measurements for HPr (black) and 2,6-$F_2$Y HPr (red) at 600 MHz.

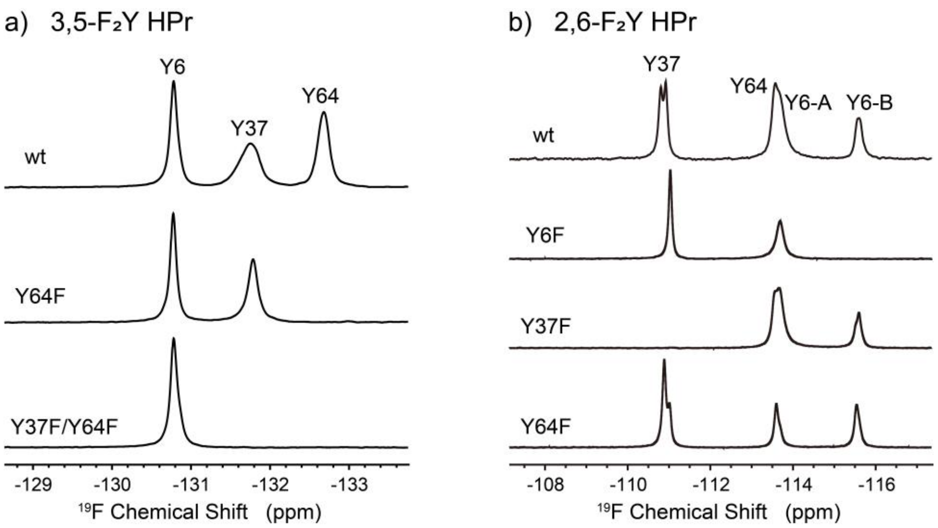

**Figure S8** $^{19}F$ peak assignments for 3,5-/2,6-$F_2Y$ HPr. a) 3,5-$F_2Y$ HPr and its Y to F variants. b) 2,6-$F_2Y$ HPr and its Y to F variants.

a) 3,5-F₂Y GB1, Y3

b) 3,5-F₂Y HPr, Y6

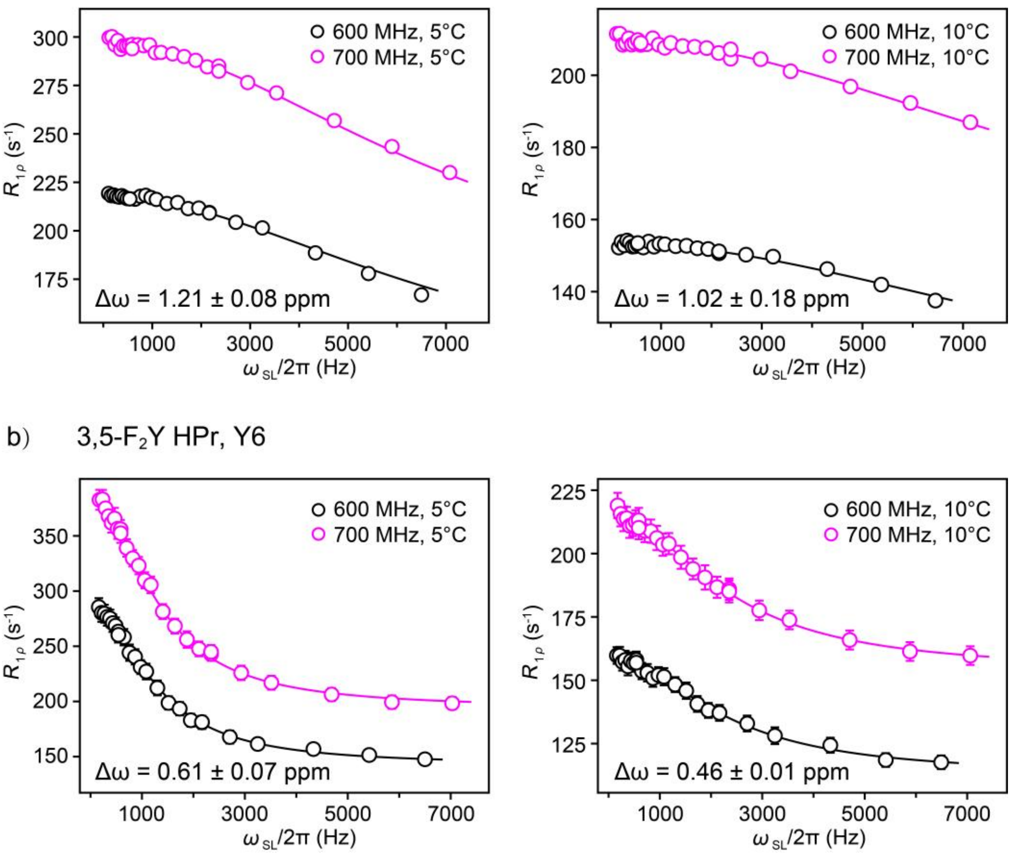

**Figure S9.** The [19]F $R_{1\rho}$ relaxation dispersion data recorded at two magnetic fields (14.1 T and 16.4 T) were fitted simultaneously, with best fits (solid lines) shown as a function of the applied spin-lock field $\omega_{SL}/2\pi$ for a) Y3 in 3,5-$F_2$Y GB1 and b) Y6 in 3,5-$F_2$Y HPr.

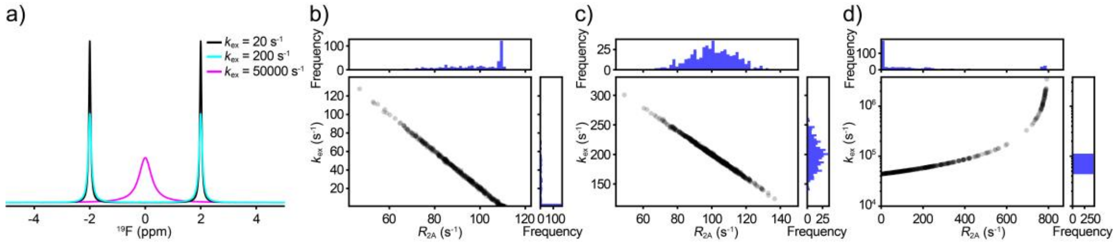

**Figure S10.** Correlation analysis between the exchange rate ($k_{ex}$) and the intrinsic transverse relaxation rate in the absence of exchange ($R_{2,0}$). a) Simulated $^{19}$F spectra with $p_1 = p_2 = 0.5$, $\Delta\omega_{F,AB}$ = 4 ppm at 500 Mhz, $R_{2,0}$ = 100 s$^{-1}$, with $k_{ex}$ = 20 s$^{-1}$ (black), 200 s$^{-1}$ (cyan) and 50000 s$^{-1}$ (magenta), respectively. (b–d) Correlation plots of fitted $R_{2A,0}$ versus $k_{ex}$ in the b) very slow exchange regime ($k_{ex}$ = 20 s$^{-1}$; $k_{ex} \ll R_{2,0}, \Delta\omega_{F,AB}$), c) slow exchange regime ($k_{ex}$ = 200 s$^{-1}$; $R_{2,0} < k_{ex} \ll \Delta\omega_{F,AB}$), and d) fast exchange regime ($k_{ex}$ = 50000 s$^{-1}$; $k_{ex} \gg \Delta\omega_{F,AB}$) from 500 Monte Carlo simulations with 1% spectral noise.

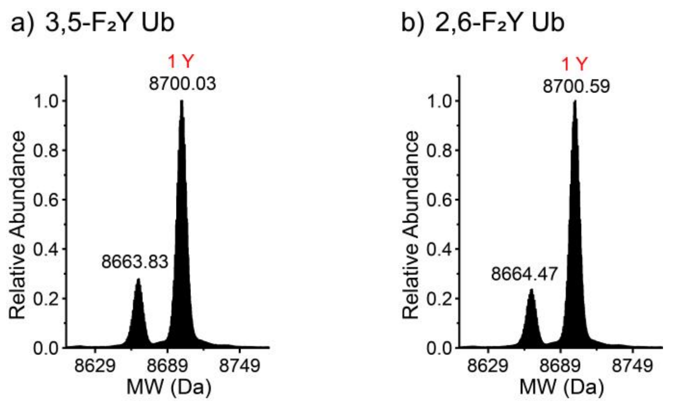

**Figure S11.** Mass spectra of a) 3,5-$F_2$Y Ub and b) 2,6-$F_2$Y Ub. The red marks indicate the number of tyrosine residue successfully labeled with di-fluorotyrosine. The yield of proteins with fluorine substitution are 78.4% and 80.7%, respectively.

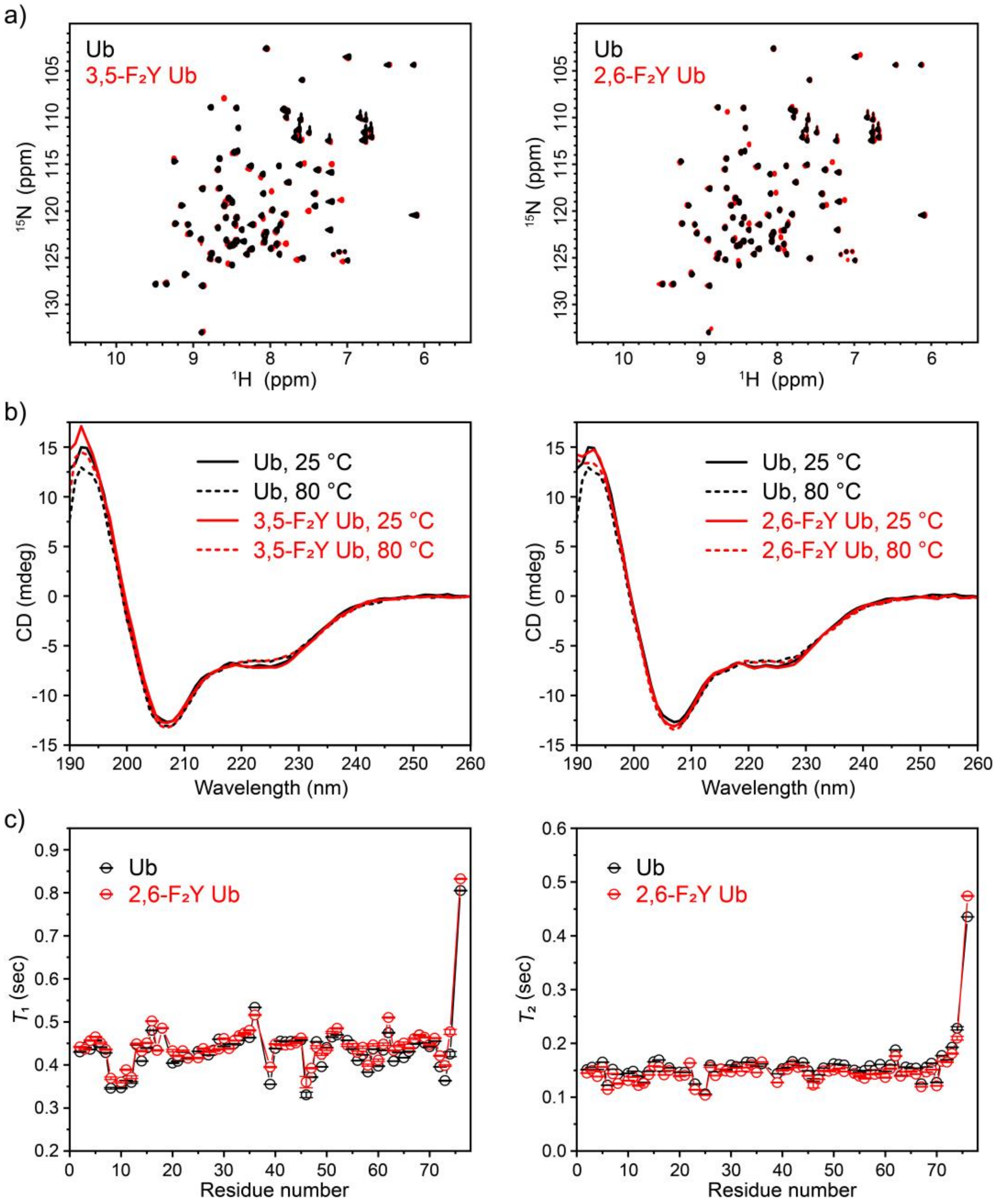

**Figure S12.** $^1$H-$^{15}$N HSQC spectra, CD spectra and $^{15}$N relaxation data for $F_2$Y Ub. a) Overlaid $^1$H-$^{15}$N HSQC spectra of Ub (black) and 3,5-$F_2$Y Ub or 2,6-$F_2$Y Ub (red). b) Overlaid CD spectra of Ub (black) and 3,5-$F_2$Y Ub or 2,6-$F_2$Y Ub (red) at 25 °C (solid) and 80 °C (dashed), respectively. c) $^{15}$N $T_1$ and $T_2$ measurements for Ub (black) and 2,6-$F_2$Y Ub (red) at 600 MHz.

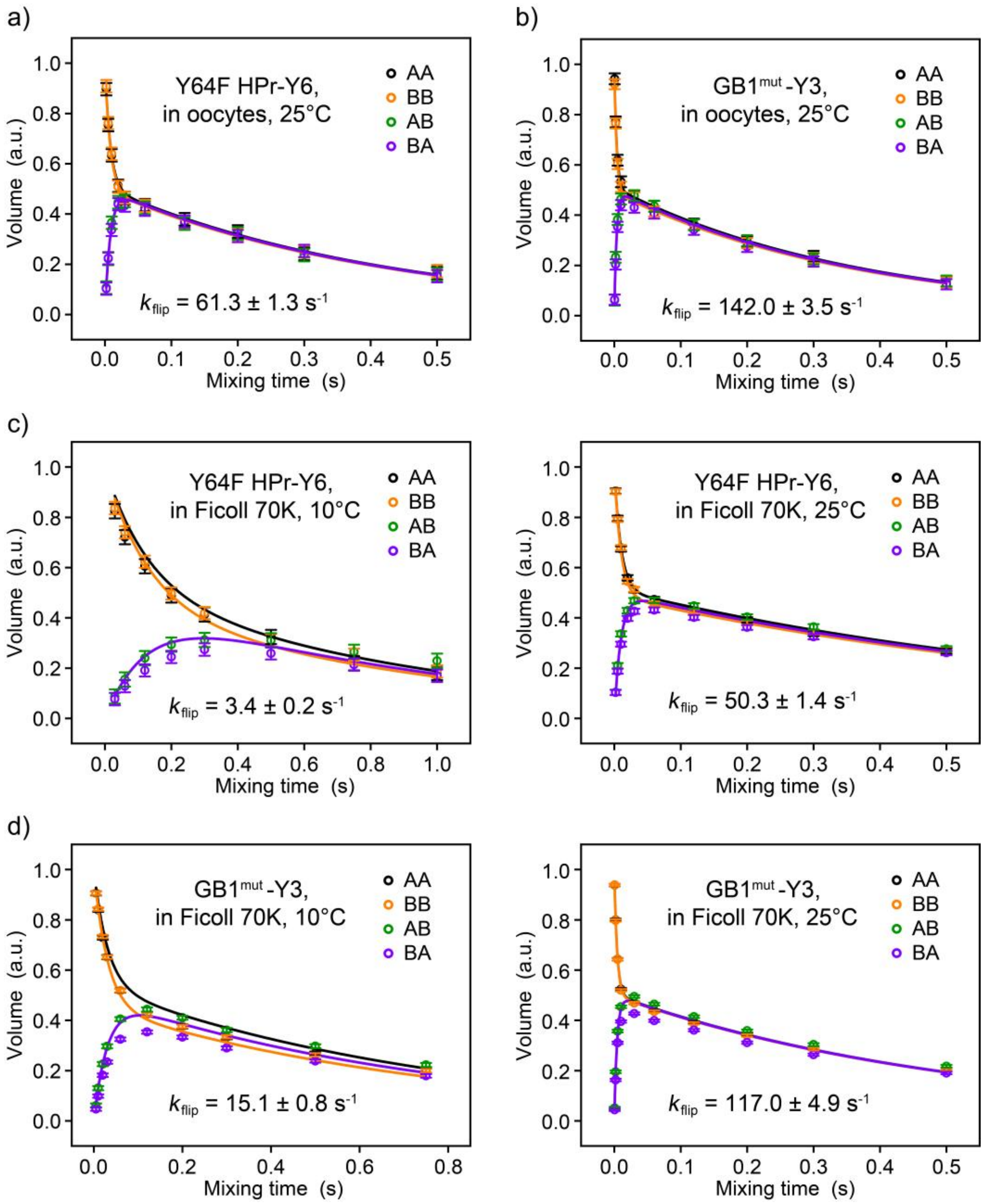

**Figure S13.** $^{19}$F-$^{19}$F EXSY experiments for measuring the ring-flip rates in *X. laevis* oocytes and in Ficoll 70K. The solid lines are best fits of the auto (AA and BB) and exchange (AB and BA) peak volumes at various mixing times to the eqs. 4-7 in the Supporting Information for a) Y6 of 2,6-$F_2$Y Y64F HPr and b) Y3 of 2,6-$F_2$Y GB1$^{mut}$ in *X. laevis* oocytes at 25 °C, and for c) Y6 of 2,6-$F_2$Y Y64F HPr and d) Y3 of 2,6-$F_2$Y GB1$^{mut}$ in 300 g/L Ficoll 70K at 10 °C (left) and 25 °C (right).

**Table S1.** $^{19}$F $R_2$ ($s^{-1}$) values of Y3 in GB1$^{mut}$ and Y6 in HPr$^{mut}$ measured at 500 MHz.

| T (°C) | 2,6-$F_2$Y Y33F/Y45F GB1 | | 2,6-$F_2$Y Y37F/Y64F HPr | |
|---|---|---|---|---|
| | Y3-A | Y3-B | Y6-A | Y6-B |
| -10 | | | 281.9 ± 7.9 | 242.4 ± 3.2 |
| 0 | 135.6 ± 1.9 | 131.0 ± 6.3 | 198.0 ± 2.6 | 169.8 ± 1.9 |
| 10 | 120.6 ± 0.7 | 110.4 ± 4.4 | 143.1 ± 1.2 | 127.5 ± 1.5 |

**Table S2.** $k_{flip}$ ($10^3$ $s^{-1}$) values for 3,5-$F_2$Y and non-fluorinated GB1 and HPr.

| T (°C) | GB1, Y3 | | GB1, Y45 | | HPr, Y6 | |
|---|---|---|---|---|---|---|
| | $^{19}$F $R_{1\rho}$ RD | $^{1}$H $R_{1\rho}$ RD | $^{19}$F $R_{1\rho}$ RD | $^{13}$C $R_{1\rho}$ RD | $^{19}$F $R_{1\rho}$ RD | $^{1}$H LS analysis |
| 35.0 | 290[a] | 46 ± 3[b] | | | 58.5[a] | 84.0[c] |
| 30.0 | 195[a] | 26 ± 2[b] | 77 ± 7 | 31 ± 6[b] | 39.5[a] | 53.9[c] |
| 25.0 | 129[a] | 15 ± 2[b] | 51 ± 5 | 11 ± 2[b] | 26.4[a] | 27.0[c] |
| 20.0 | | | 34 ± 3 | 6 ± 2[b] | 17.3[a] | 16.7[c] |
| 15.0 | | | | | 10.6 ± 0.8 | 7.4[c] |
| 12.5 | 43 ± 2 | | | | 9.2 ± 0.3 | |
| 10.0 | 35 ± 2 | | 14[a] | | 7.4 ± 0.3 | 4.3[c] |
| 7.5 | 28 ± 2 | | | | 5.5 ± 0.5 | |
| 5.0 | 22 ± 1 | | | | 4.1 ± 1.0 | |

[a] Extrapolated from the fitted *ΔH* and *ΔS* using the Eyring equations.

[b] Taken from the work of M. Dreydoppel et al.[29-30]

[c] Taken from the work of M. Hattori et al.[31]

**Table S3.** $k_{flip}$ ($s^{-1}$) values for 2,6-$F_2$Y labeled proteins determined by $^{19}F$-$^{19}F$ EXSY.

| T (°C) | Y33F/Y45F GB1, Y3 | | | HPr, Y6 | Y64F HPr, Y6 | | |
|---|---|---|---|---|---|---|---|
| | in buffer | in oocyte | in Ficoll[a] | in buffer | in buffer | in oocyte | in Ficoll[a] |
| 10 | 20.1 ± 0.5 | 19.2 ± 0.7 | 15.1 ± 0.8 | 3.4 ± 0.2 | 3.4 ± 0.1 | 3.9 ± 0.2 | 3.4 ± 0.2 |
| 25 | 157.1 ± 4.4 | 142.0 ± 3.5 | 117.0 ± 4.9 | 63.5 ± 4.0 | 61.2 ± 1.7 | 61.3 ± 1.3 | 50.3 ± 1.4 |

[a] 300 g/L Ficoll 70K.
